# Fine tuning energy metabolism in skeletal muscle: Discovery of a novel autoinhibitory mechanism in the N-terminal extension of AMPKγ3

**DOI:** 10.64898/2026.08.16.744724

**Authors:** Ashley J. Ovens, Muhammad N.H. Khabib, Dingyi Yu, Naomi X.Y. Ling, William J. Smiles, Ashfaqul Hoque, Danise Ann Onda, Andrea C. Poblete Goycoolea, Meiqi Cao, Geoff X.Y. Zhang, Benjamin R. Turner, Larissa Doughty, Ching-Seng Ang, Christopher R. Horne, John W. Scott, Kei Sakamoto, Michael W Parker, Bruce E. Kemp, Sandra Galic, Jonathan S. Oakhill, Christopher G. Langendorf

**Affiliations:** Protein Engineering in Immunity & Metabolism, St. Vincent’s Institute of Medical Research, Fitzroy, VIC, 3065, Australia; Metabolic Signalling Laboratory, St. Vincent’s Institute of Medical Research, Fitzroy, VIC, 3065, Australia; Protein Chemistry & Metabolism, St. Vincent’s Institute of Medical Research, Fitzroy, VIC, 3065, Australia; Diabetes & Metabolic Disease, St. Vincent’s Institute of Medical Research, Fitzroy, VIC, 3065, Australia; ACRF Rational Drug Discovery Centre, St. Vincent’s Institute of Medical Research, Fitzroy, VIC, 3065, Australia; Structural Biology and Computational Design Laboratory, Department of Biochemistry and Pharmacology, Bio21 Molecular Science and Biotechnology Institute, University of Melbourne, Parkville, VIC, 3052, Australia; Mass Spectrometry and Proteomics Facility, Bio21 Molecular Science and Biotechnology Institute, University of Melbourne, Parkville, VIC, 3052, Australia; Walter and Eliza Hall Institute of Medical Research, Parkville, VIC, 3052, Australia; Drug Discovery Biology, Monash Institute of Pharmaceutical Sciences, Monash University, Parkville, VIC, 3052, Australia; Novo Nordisk Foundation Center for Basic Metabolic Research, University of Copenhagen, Copenhagen, 2200, Denmark; Faculty of Health Sciences, Australian Catholic University, Fitzroy, VIC, 3065. Australia; Department of Medicine, University of Melbourne, Parkville, VIC, 3010, Australia; Research Program for Receptor Biochemistry and Tumour Metabolism, University Hospital of the Paracelsus Medical University, Salzburg, 5020, Austria

**Keywords:** AMP-activated protein kinase (AMPK), skeletal muscle physiology, enzyme kinetics, kinase, allosteric modulation, diabetes, metabolic disease

## Abstract

AMP-activated protein kinase (AMPK) regulates metabolism in response to metabolic stress that includes stimulating glucose uptake in skeletal muscle independently of the canonical insulin signalling pathway, positioning it as an attractive therapeutic target for insulin resistance and type 2 diabetes mellitus (T2DM). AMPK is an αβγ heterotrimer, with multiple isoforms for each subunit enabling the formation of 12 different complexes with distinct tissue expression profiles. Among these, the α2β2γ3 complex is predominantly expressed in skeletal muscle, the major site of glucose disposal and a highly desirable therapeutic target for T2DM. Here, we characterise the functional role of a unique, 182 residue N-terminal extension (NTE) within γ3 subunit. Deletion of the γ3-NTE from α2β2γ3 complex increases basal AMPK activity without affecting activation by AMP or pharmacological AMPK activators, demonstrating the γ3-NTE performs an autoinhibitory function. Using complementary biophysical techniques, including hydrogen-deuterium exchange-mass spectrometry, surface plasmon resonance, chemical crosslinking and co-pulldowns, we identified a 39-residue sequence in the γ3-NTE (residues 129-168), that directly interacts with the αC-helix of the AMPK kinase domain small lobe, a key regulatory element in many protein kinases. Using AlphaFold3, we probe the interaction predicted to take place between a γ3-NTE α-helix (γ3-iHelix; ∼T142-E154) and the αC-helix in the α2β2γ3 complex. These findings provide the groundwork for developing novel T2DM therapies that target AMPK activation selectively in skeletal muscle involving reversal of the γ3 autoinhibition.

## Introduction

Every cellular biological process depends on sufficient cellular energy, whereby disruptions to the normal homeostatic control of energy balance can lead to a variety of pathophysiological changes that contribute to the progression of metabolic disease (1, 2). To maintain metabolic homeostasis, the balance of cellular energy is regulated by the AMP-activated protein kinase (AMPK) via control of a range of fundamental bioenergetic processes (e.g., protein turnover, glycolysis, lipid oxidation etc.) (3, 4). AMPK is considered a promising therapeutic target for metabolic diseases including type 2 diabetes mellitus (T2DM), due to its role in stimulating insulin-independent glucose uptake in skeletal muscle (5–7). AMPK is a phylogenetically conserved serine protein kinase that forms an αβγ heterotrimeric complex. Multiple isoforms for each AMPK subunit exist in mammals (α1/α2, β1/β2, γ1/γ2/γ3), giving rise to 12 distinct heterotrimeric combinations with unique biochemical profiles (8). The expression profiles of each isoform vary significantly, ranging from ubiquitous to highly distinct, tissue-specific distribution. For example, α1, β1, γ1 and γ2 are ubiquitously expressed, with γ2 playing an important functional role in the heart (9). Conversely, α2, β2 and γ3 are enriched in skeletal muscle (10–12), with γ3 displaying the most restricted expression profile being largely confined to glycolytic/fast-twitch muscle fibres (e.g., extensor digitorum longus; EDL) (13, 14), and to a lesser extent brown adipose tissue (15, 16). Immunoprecipitation of AMPK heterotrimers indicates that only α1β2γ1, α2β2γ1 and α2β2γ3 complexes can form in human skeletal muscle (10, 17), with γ3 complexes comprising only 20% of the pool of AMPK heterotrimers in glycolytic/fast-twitch skeletal muscle (18).

The α-subunit contains the canonical serine/threonine kinase domain (α-KD) and regulatory elements, including an autoinhibitory domain (AID), interacting motifs, and a largely disordered serine/threonine-rich loop (ST-loop) which contains known regulatory phosphorylation sites. (19). The regulatory β-subunit contains a carbohydrate binding module (β-CBM), an α-γ-subunit binding sequence (α-γ-SBS) and is myristoylated on β1/2-G2, which is required for interaction with cellular membranes (e.g., lysosomes, mitochondria)(19–22). Lastly, the regulatory γ-subunit contains four tandem cystathionine-β-synthase (CBS) repeats that form the energy-sensing AMP-, ADP- and ATP-binding sites; site 2 is regarded as being unoccupied, site 4 thought to permanently house AMP, and sites 1 and 3 interchangeably binding to the three nucleotides (19, 23). Preceding this region is a short pre-CBS sequence that interacts with the β-subunit to stabilise the heterotrimeric complex (24).

The canonical mechanism of AMPK activation involves phosphorylation of the α-KD activation loop on α1/2-T174/172 (hereafter pT172) by the upstream kinases liver kinase B1 (LKB1) and Ca^2+^-calmodulin dependent kinase kinase-2 (CAMKK2) (25–29). pT172 is important for stabilising the αC-helix of the α-KD in an active, inward conformation that favourably aligns catalytic residues in the core of the enzyme (30). Additionally, AMP or ADP binding to CBS sites in the γ-subunit induce conformational changes that both promote α−T172 phosphorylation and shield the phosphorylated residue from phosphatases, whereas AMP alone can allosterically activate the complex roughly 2-fold (31, 32). Additional allosteric regulation of AMPK activity is achieved through the allosteric drug and metabolite (ADaM) site (33, 34), a hydrophobic cleft formed at the interface of the α-KD N-lobe and β-CBM (33). Endogenous nutrient metabolites such as palmitoyl-CoA as well as synthetic small-molecules (e.g., SC4, MSG011, MK8722, PF739, 991) bind to the ADaM site to stimulate AMPK activity (5, 6, 35, 36). Mechanistically, ADaM site ligands stabilize the α-KD and β-CBM interface, which positions an α-helix that immediately follows on from the β-CBM (called the C-interacting helix) to pack against and stabilises the αC-helix of the α-KD, promoting an optimal catalytic conformation activating AMPK independent of α−T172 phosphorylation (30, 34). AMPK is also regulated by other phosphorylation sites (37), particularly phosphorylation of the α-ST-loop by members of the AGC family of protein kinases (e.g., Akt) (38). One phosphorylation site on the β-subunit, S108, is situated within the ADaM site and sensitises AMPK to allosteric ligands (34, 39).

Studies in rodent and non-human primate models have demonstrated that direct pan-AMPK-activating drugs induce sustained improvements in glucose homeostasis through activation of skeletal muscle AMPK, including reductions in hyperglycaemia and insulin resistance without causing hypoglycaemia (5, 6). However, systemic activation by the pan-AMPK activator MK8722 was shown to induce side-effects such as cardiac hypertrophy in non-human primates and mice, highlighting the need for more targeted, isoform-selective therapeutic approaches. Although AMPKγ3 is not required to maintain normal blood glucose homeostasis, it remains a promising therapeutic target to pharmacologically modulate glucose uptake in skeletal muscle. We recently showed that AMPKγ3 is absolutely required for skeletal muscle glucose uptake via AMPK activation by 5-aminoimidazole-4-carboxyamide ribonucleotide (AICAR; a prodrug of the AMP mimetic ZMP) (40), while Kido and colleagues revealed AMPKγ3 regulates muscle glucose clearance during post-exercise recovery (41).

The AMPK γ2 and γ3 isoforms each contain a distinct N-terminal extension (NTE) that precedes the CBS motifs, consisting of 259 and 182 amino acid residues, respectively (42). The γ2-NTE contains a nuclear localisation sequence (43), an undefined region that protects pT172 from phosphatases (44, 45), and multiple phosphorylation sites of unknown function (46). Recent evidence highlights the importance of γ3 isoform in metabolic disease with a phosphorylation site, γ3-S65, unique to the human γ3-NTE which is strongly associated with insulin resistance (47). The functional role of the γ3-NTE is largely undefined, although previous studies provide some insight into the general function of γ3-AMPK. γ3-AMPK complexes are largely resistant to AMP allosteric activation (45, 48); but are sensitive to AMP- and ADP-mediated protection against pT172 dephosphorylation (45). The γ1- and γ3-subunit isoforms were reported to comparably bind adenine nucleotides (AMP/ADP/ATP) and the AMP-mimetic ZMP with comparable affinity (49). However, there is also evidence that AMP binds more strongly to site 1 in γ3 compared to γ1 (48). It was reported that ATP affinity in the active site is between four- to 10-fold higher for γ3 complexes. Since the CBS binding sites are highly conserved between γ1 and γ3 and located ∼50 Å away from the ATP binding pocket within the α-KD, it is plausible the γ3-NTE may be interacting with the α-KD to cause this reduction in ATP affinity (49).

In this study, we investigated the functional role of the γ3-NTE and demonstrated using multiple biophysical approaches that the γ3-NTE contains an autoinhibitory sequence that directly interacts with the α-KD to suppress kinase activity. This autoinhibitory sequence is predicted to form a helical structure we suspect inhibits AMPK activity through misalignment of the αC-helix within the α-KD. Our results thus provide new insights into the molecular mechanisms regulating AMPK complexes containing the γ3 isoform.

## Results

### The γ3-NTE contains an autoinhibitory sequence

To investigate regulatory role for the γ3-NTE, we expressed all human α/β isoform combinations with C-terminal FLAG-fusion wild type γ3 (γ3-WT) or NTE-deleted γ3 (γ3-ΔNTE) lacking residues 1-168 from HEK293T/17 cells, followed by FLAG affinity purification (Fig. 1A). This deletion was selected to retain the sequence of the scaffolding region (residues I183-M197) required for formation of stable AMPK heterotrimer complexes, which we confirmed by the presence of α- and β-subunits after FLAG purification of the γ3-subunit (Fig. 1A). Removal of the γ3-NTE otherwise had no effect on basal pT172, except for a ∼40% reduction in the α2β1γ3 complex (Fig. 1A, B). Despite no overt differences in the activation loop T172 phosphorylation, all γ3-ΔNTE complexes displayed significantly elevated basal activity compared to their wild-type (WT) counterparts when assayed at 200 μM ATP. The increase in activity ranged from 2.8-fold for α2β2γ3ΔNTE to 5.8-fold for α1β1γ3ΔNTE (Fig. 1C). Treatment of α2β2γ3 and α2β2γ3ΔNTE with CAMKK2 during the FLAG purification to maximally phosphorylate α-T172 increased overall basal activity, however the γ3-ΔNTE complex retained its elevated activity compared to the WT complex, indicating the increased kinase activity caused by loss of the γ3-NTE is indeed independent of pT172 (Figs. 1D & S1A, B). We also detected a 2.4-fold higher basal activity for α2β2γ3ΔNTE compared to WT when expressed in COS7 cells, demonstrating preservation of the regulatory effect of the γ3-NTE overexpression across different cell lines (Figs. 1E & S1C). Phosphorylation of acetyl-CoA carboxylase (ACC), an endogenous AMPK substrate, was significantly elevated by 2-fold in HEK293T cells expressing α2β2γ3ΔNTE compared to cells expressing WT α2β2γ3, confirming increased AMPK signalling with the γ3-NTE deletion (Fig. 1F, G). Removal of the γ3-NTE had no impact on allosteric activation of AMPK complexes by MSG011, a pan-AMPK ADaM-site activator (50), or AMP (Fig. S1D, E), suggesting γ3-ΔNTE complexes form heterotrimers functionally responsive to ligand stimulation.

**Figure 1.**
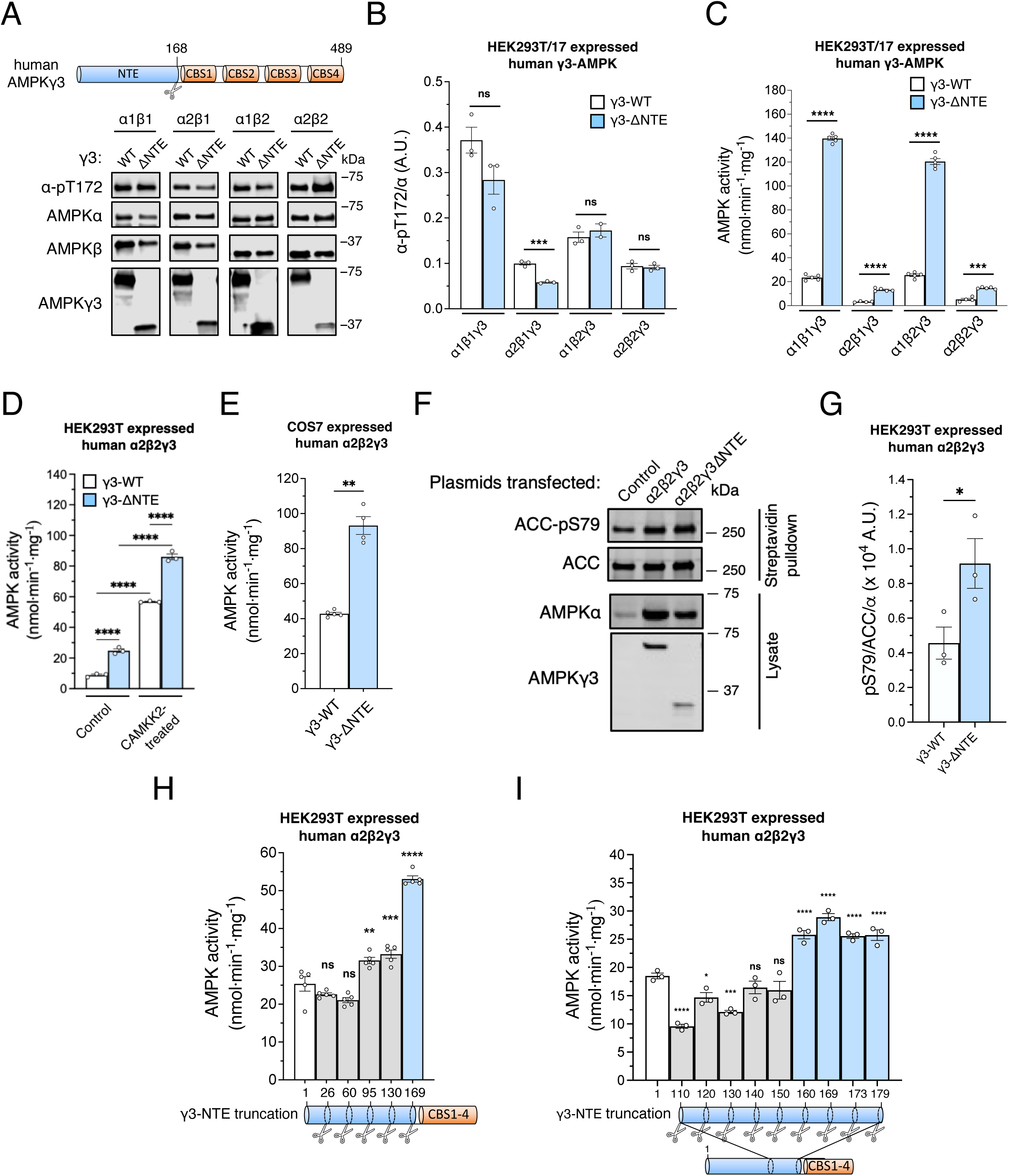
Human AMPKγ3 complexes expressed in mammalian cells and basal kinase activity. (A) Immunoblot images of α-pT172 and AMPK subunits from complexes containing γ3 wild-type (γ3-WT) or N-terminal extension deleted (γ3-ΔNTE) expressed and FLAG-purified from HEK293T/17 cells. (B) Densitometry analysis of (A) comparing basal α-pT172 between γ3-complexes. Data are shown as mean α-pT172/α (arbitrary units, A.U.) ± s.e.m.; *n* = 3. (C) Basal activities of the AMPK complexes FLAG-purified in (A). Data are shown as mean AMPK activity (nmol·min^−1^·mg^−1^) ± s.e.m.; *n* = 4/5. (D) Basal activities of AMPK α2β2-complexes containing γ3-WT or γ3-ΔNTE untreated and CAMKK2-treated during FLAG-purification from HEK293T cells. Data are shown as mean AMPK activity ± s.e.m.; *n* = 3. (E) Basal activities of the AMPK complexes FLAG-purified from COS7 cells. Data are shown as mean AMPK activity ± s.e.m.; *n* = 3. (F) Immunoblot images of total and phospho-acetyl-CoA carboxylase (ACC, ACC-pS79) Streptavidin-purified from HEK293T cells overexpressing AMPK α2β2-complexes containing γ3-WT or γ3-NTE. (G) Densitometry analysis of (F) comparing ACC-pS79 level between γ3-WT and γ3-ΔNTE complexes. pS79/ACC ratios were normalized to total AMPKα from respective lysates to account for lower expression in the γ3-ΔNTE complexes. Data are shown as mean pS79/ACC/α (A.U.) ± s.e.m.; *n* = 3. (H-I) Basal activities of AMPK α2β2-complexes containing γ3-WT or sequentially truncated γ3, as indicated. (B, C, E, & G) Statistical analyses were performed by Welch’s *t*-test vs. respective γ3-WT complexes. ns, *p* > 0.05, \**p* < 0.05, \*\*\**p* < 0.001, \*\*\*\**p* < 0.0001. (D, H & I) Statistical analyses were performed by one-way ANOVA with Dunnett’s multiple-comparisons test vs. γ3-WT. ns, *p* > 0.05, *\*p <* 0.05, \*\**p* < 0.01, \*\*\**p* < 0.001, \*\*\*\**p* < 0.0001. (A, F) Representative immunoblot images of 3 independent experiments are shown.

Following reports that γ3-AMPK has a lower affinity for ATP than γ1-AMPK (48), we sought to corroborate this observation and determine if the γ3-NTE is responsible for these differences. We used enzyme kinetic assays to compare the *K*_M_ values of different AMPK complexes. *K*_M_ values for ATP with α1β1γ3 expressed in HEK293T/17 cells were 3.5-fold and 1.8-fold higher than α1β1γ1 and α1β1γ2, respectively (Fig. S2A), whereas the *K*_M_ value for ATP with α2β2γ3 expressed in bacteria was 8.9-fold higher than α2β2γ1 (Fig. S2B). These findings are consistent with a previous report demonstrating 10-fold higher *K*_M_ values for ATP with bacterial-expressed α2β2γ3 compared to α2β2γ1, leading to the hypothesis that the γ3-NTE either directly, or indirectly, impairs ATP binding at the kinase domain active site (48). We determined the *K*_M_ values for ATP for γ3-WT and γ3-ΔNTE complexed to each α/β isoform combination (Table 1; Fig. S2C-F). Deletion of the γ3-NTE reduced *K*_M_ values for ATP by 1.1-, 3.8-, and 1.3-fold in α2β1γ3, α1β2γ3, and α2β2γ3 complexes, respectively, but the ATP *K*_M_ value increased ∼2-fold in α1β1γ3 complex (Table 1). Specific activities of all γ3-ΔNTE complexes were markedly higher than their respective WT complexes at all assayed concentrations of ATP, with the maximal enzyme velocity (V_max_) increasing ∼7-fold for both β1 complexes and 2- to 3-fold for β2 complexes. Consequently, all γ3-ΔNTE complexes exhibited higher catalytic efficiency (V_max_/*K*_M_) than their respective WT complexes, showing a 3.6-fold increase in α1β1 and α2β2 complexes, and 7.8-fold in α1β2 and α2β1 complexes (Table 1). Combined, these data indicate that the γ3-NTE negatively regulates AMPK activity independently of pT172 across all α/β isoform compositions, with the suppressive effect partially explained by γ3-NTE-mediated interference of ATP binding the active site.

**Table 1.** Kinetics of AMPK complexes containing γ3-WT or γ3-ΔNTE under varying ATP concentrations. Values were derived using non-linear regression analysis fitted to the Michaelis-Menten equation of AMPK activity assay across varying ATP concentrations. Catalytic efficiency values were calculated as the quotient of *V_max_*/*K_M_*. Each parameter is presented as value ± s.e.m.

| AMPK complex | $V_{\max}$<br>(nmol·min<sup>-1</sup>·mg<sup>-1</sup>) | ATP $K_M$ (μM) | Catalytic efficiency<br>(nmol·min<sup>-1</sup>·mg<sup>-1</sup>·μM<sup>-1</sup>) |
| --- | --- | --- | --- |
| α1β1γ3 | 48.0 ± 1.7 | 90.6 ± 13.2 | 0.53 ± 0.06 |
| α1β1γ3ΔNTE | 325.8 ± 10.4 | 172.6 ± 18.2 | 1.89 ± 0.15 |
| α2β1γ3 | 4.3 ± 0.4 | 234.6 ± 60.1 | 0.018± 0.003 |
| α2β1γ3ΔNTE | 30.0 ± 1.9 | 208.5 ± 41.1 | 0.14 ± 0.02 |
| α1β2γ3 | 83.2 ± 10.3 | 440.9 ± 134.0 | 0.19 ± 0.04 |
| α1β2γ3ΔNTE | 169.3 ± 7.7 | 113.3 ± 19.7 | 1.49 ± 0.21 |
| α2β2γ3 | 7.8 ± 0.5 | 161.2 ± 38.9 | 0.05 ± 0.01 |
| α2β2γ3ΔNTE | 22.8 ± 0.7 | 123.9 ± 15.2 | 0.184 ± 0.02 |

To identify a putative autoinhibitory sequence within the γ3-NTE, we expressed and purified sequential truncations of γ3-NTE co-transfected with α2 and β2 in HEK293T cells (Figs. 1H & S3A). All truncated γ3-NTE constructs formed stable and active heterotrimeric complexes (Fig. S3A), demonstrating comparable MSG011 and AMP activation profiles to those of γ3-WT (Fig. S3B). N-terminally truncated constructs beginning at residues γ3-M26 or γ3-W60 produced no measurable change in basal AMPK activity compared to γ3-WT, suggesting the autoinhibitory region lies further downstream in the primary sequence. Subsequent N-terminal truncated constructs beginning at residues γ3-K95 or γ3-T130 resulted in 1.3-fold increases in basal activities versus γ3-WT, suggesting the region between γ3-W60 and γ3-P94 plays a minor role in autoinhibition. Full release of autoinhibition was only achieved with the γ3-ΔNTE construct (truncation to γ3-P168; Fig. 1H), indicating a region in γ3 between T130 and P168 is primarily responsible for γ3-NTE-mediated AMPK inhibition.

To pinpoint the specific inhibitory sequence within this region of γ3, we generated an additional series of N-terminal truncations at intervals of 4-10 residues between P110 and P179 and co-expressed and purified these constructs with α2 and β2 as stable complexes from HEK293T cells (Fig. S3C). Truncating the γ3-NTE up to γ3-M185 impaired AMPK complex formation as indicated by low levels of the α- and β-subunit co-purifying with the γ3-subunit (Fig. S3C), presumably due to deletion of two amino acids from the scaffolding region shown to be required for AMPK complex formation (24, 42). N-terminal deleted constructs beginning between residues γ3-E110 and γ3-E150 resulted in either unchanged or reduced activity compared to WT (Fig. 1I), with reduced activity possibly being accounted for by significantly lower pT172 levels (Fig. S3C, D). In this set of protein preparations, truncations up to γ3-T130 did not produce the 2.8-fold increase in AMPK activity observed in independent experiments (Fig. 1E), perhaps reflecting the high sensitivity of this specific construct to subtle changes in protein stability or buffer composition. However, truncated constructs beginning at residues γ3-E160 and beyond resulted in enhanced AMPK activity compared to WT (Fig. 1I), with modestly elevated pT172 associated only with the construct beginning at residue γ3-C160 (1.2-fold vs. WT) (Fig. S3D). These findings demonstrate that the autoinhibitory function of the γ3-NTE is largely mediated by a short stretch of amino acids between γ3-E150 and γ3-L159 (EGLLEERPAL).

### Phosphorylation profiling of γ3-NTE deleted AMPK complexes

Using a combination of immunoblotting and quantitative MS-based analyses (46), we investigated the phosphorylation profiles of γ3-ΔNTE complexes compared with γ3-WT across all α/β isoform combinations. There were no differences in phosphorylation of the inhibitory ST-loop sites α1/2-S487/491 by immunoblot, except for α2β1γ3, which showed a reduction on γ3-ΔNTE compared to γ3-WT complexes (Fig. 2A, B), which is despite α2-S491 being a well-known autophosphorylation site (37) and expected to increase commensurate with kinase activity. α1-S487 was also quantified by MS and displayed no differences in phosphorylation between γ3-ΔNTE and γ3-WT complexes, consistent with immunoblot data (Fig. 2C). Changes in β2-pS108 autophosphorylation, known to activate AMPK (37), was quantified by MS; both αβ2γ3ΔNTE complexes showed increased β2-pS108 compared to WT, suggesting elevated *in cellulo* AMPK activity (Fig. 2D). mTORC1 substrates α1/2-S347/345 (inhibitory) and α2-S377 (function unknown) (51, 52) displayed reduced phosphorylation levels in γ3-ΔNTE complexes (Fig. 2E, F). Our MS analysis also included ST-loop phosphosites α1/2-S477/481 (unknown effect) and α1/2-T481/485 (inhibitory), both substrates for GSK3β (53). α1-pS477 was unchanged, whereas α2-pS481 was lower on both γ3-ΔNTE complexes (Fig. S4A), and α1/2-pT481/485 was lower except in the α2β2 complex (Fig. S4B). Additional uncharacterised ST-loop phosphorylation sites analysed by MS included α1-pS499, which was reduced on γ3-ΔNTE complexes (Fig. S4C), and α2-S501 that remained unchanged compared to γ3-WT (Fig. S4D). β1/2-S182/184 is another validated inhibitory mTORC1 substrate that is highly resistant to dephosphorylation (46), likely explaining why its levels remained the same on γ3-ΔNTE complexes (Fig. S4E). This analysis reveals that γ3-ΔNTE complexes exhibit elevated *in cellulo* AMPK activity, as evidenced by increased autophosphorylation of β2-S108. This heightened activity is associated with a concomitant reduction in the phosphorylation of mTORC1 and GSK3β sites known to reduce AMPK activity, likely mediated through AMPK-dependent inhibitory feedback loops.

**Figure 2.**
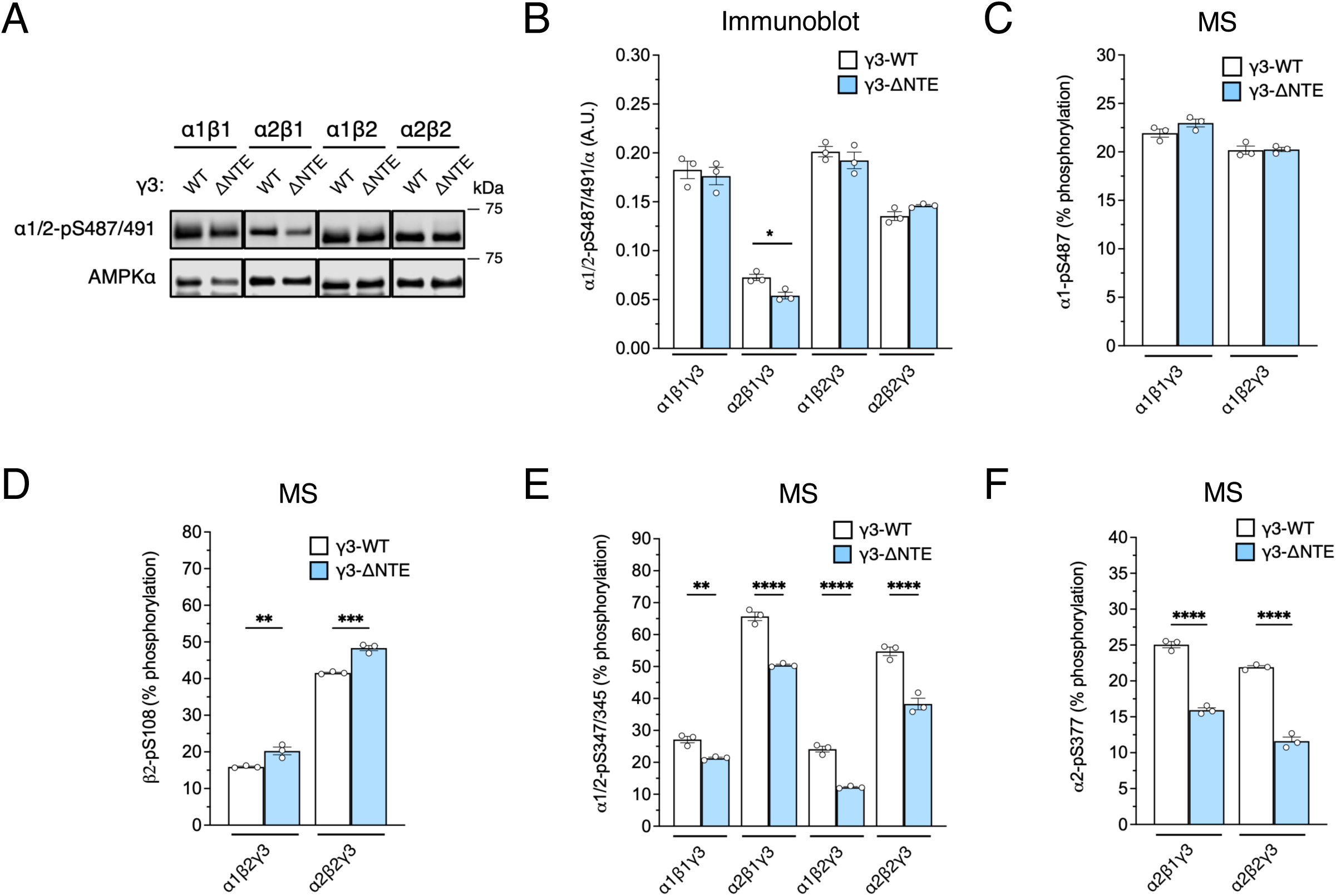
Phosphorylation site profiling of AMPK complexes containing γ3-WT or γ3-ΔNTE. (A) Immunoblot images of α1/2-pS487/491 and total AMPKα from complexes containing γ3 wild-type (γ3-WT) or N-terminal extension deleted (γ3-ΔNTE) expressed and FLAG-purified from HEK293T/17 cells. Representative immunoblot images of 3 independent experiments are shown. (B) Densitometry analysis of (A) comparing basal α1/2-pS487/491 levels between γ3-complexes, as indicated. Data are shown as mean α1/2-pS487/491/α (A.U.) ± s.e.m.; *n* = 3. Area under the curve stoichiometry analysis of LC-MS data for tryptic-digested AMPK from (A) for multiple phosphorylation sites (C) α1-pS487, (D) β2-pS108, (E) α1/2-pS347/345, and (F) α2-pS377. Data are adjusted for the flyability ratios as previously described (46), and presented as mean % phosphorylation ± s.e.m.; *n* = 3. Statistical analyses were performed by Welch’s *t*-test vs. respective γ3-WT complexes. \**p* < 0.05, \*\**p* < 0.01, \*\*\**p* < 0.001, \*\*\*\**p* < 0.0001.

### The γ3-NTE interacts directly with the α2-KD

Since the γ3-NTE contains an autoinhibitory region that restricts AMPK activity, independent of reductions in pT172, we hypothesised it may directly interact with the α-KD to allosterically modulate AMPK. To test for a direct interaction between the two proteins, we expressed and purified a γ3_2-168_ (γ3-NTE) fragment with a C-terminal AVI-tag from *E. coli* (Fig. 3A-C). This 20 kDa fragment had an expected mass confirmed by time of flight-mass spectrometry (TOF-MS; Fig. S5A), but like the full-length γ3 protein (Fig. 1A), migrated ∼10 kDa slower (at around at 30 kDa) in SDS-PAGE gels (Fig. 1A). However, mammalian cell-expressed γ3-ΔNTE displayed anticipated SDS-PAGE mobility, migrating at ∼37 kDa (Fig. 1A), suggesting the γ3-NTE is responsible for this discrepancy, analogous to the different β-subunit isoforms that have virtually identical molecular weights yet disparate migratory patterns in SDS-PAGE (54, 55). Such aberrant electrophoretic mobility is not uncommon for proteins with densely acidic regions, such as those found in the γ3-NTE (56).

**Figure 3.**
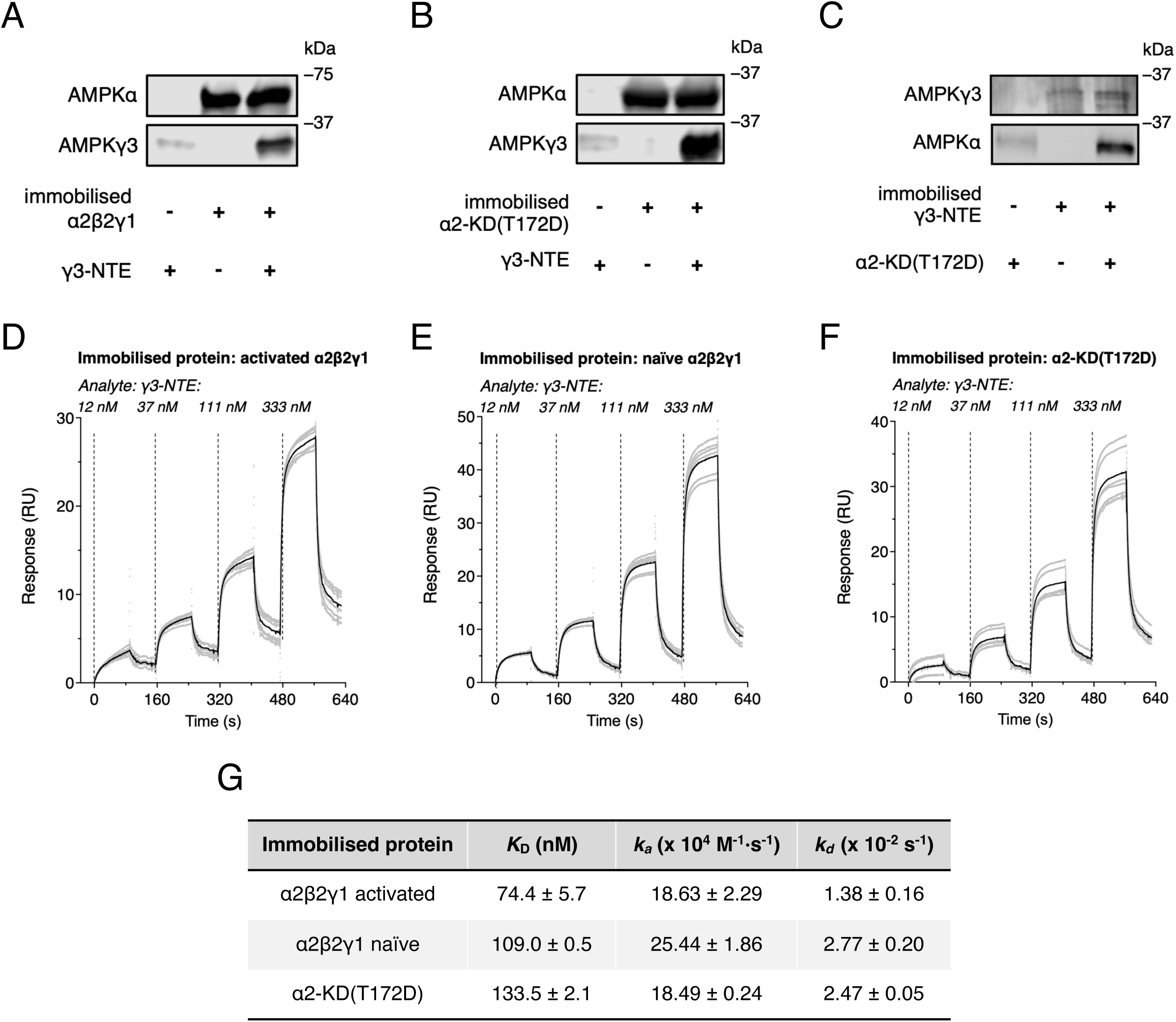
Analyses of interactions between AMPK γ3-NTE and α2-KD or α2β2γ1 by co-pulldown and surface plasmon resonance (SPR). Immunoblot images of co-pulldown experiments using Ni^2+^-NTA-immobilised (A) α2β2γ1 complex, and (B) α2-KD(T172D) to capture soluble γ3-NTE. (C) Immunoblot images of reciprocal co-pulldown experiments using Streptavidin Sepharose-immobilised γ3-NTE to capture soluble α2-KD(T172D). Representative immunoblot images of 3 independent experiments are shown. Double-referenced SPR sensorgrams against reference cell and buffer reference signal (grey dots) and their averages (black line) from three replicates (*n* = 3; each replicate contains a repeat producing a total of six sensorgrams) using immobilised (D) activated α2β2γ1, (E) naïve α2β2γ1, and (F) α2-KD(T172D) to capture soluble γ3-NTE. (G) Association (*k*_a_) and dissociation (*k*_d_) rate constants of the γ3-NTE interaction with the 3 immobilised proteins determined by global fitting of the sensorgrams to a 1:1 binding model using Biacore T200 Evaluation Software v2.0 (D-F). Equilibrium dissociation constants (*K_D_*) were derived from the calculated rate constants. Data are shown as mean for each constant ± s.d.; *n* = 3.

The purified γ3-NTE protein was utilised for *in vitro* co-pulldown and immunoblotting assays. Since γ1 does not contain a large NTE, we reasoned that any potential γ3-NTE-interacting site in the *α*-KD would be available for binding in the γ1-AMPK complexes. We did not used γ3-ΔNTE complex because its bacterial expression failed (data not shown). We detected a direct interaction between soluble γ3-NTE and activated *α*2β2*γ*1 (CAMKK2-treated) immobilised on Ni^2+^-NTA (Fig. 3A). Robust γ3-NTE enrichment was also observed using the constitutively active *α*2-KD(T172D) immobilised as the protein bait (Fig. 3B). The interaction was evident in a reciprocal pulldown experiment using biotinylated γ3-NTE immobilised on Streptavidin Sepharose as the bait and incubated with soluble *α*2-KD(T172D) (Fig. 3C). To determine whether AMPK ligands and inhibitors altered the α-KD/γ3-NTE interaction, we co-incubated with Mg^2+^-ATP, and ATP-competitive kinase inhibitors staurosporine, compound C and SBI-0206965. These experiments were performed using a kinase dead *α*2(D139A)β2γ1 mutant as bait for the γ3-NTE to avoid unintended effects of enzymatic activity. Neither the kinase inhibitors nor Mg^2+^-ATP up to a concentration of 500 µM had any observable effect on binding, however 5 mM Mg^2+^-ATP almost completely abolished the interaction independent of Mg^2+^ alone (Fig. S6A, B). We also tested AMPK peptide substrates SAMS and S108tide (39), with SAMS but not S108tide enhancing the binding of γ3-NTE (Fig. S6C). AMP concentrations above 50 µM also disrupted the interaction (Fig. S6C). These disruptive effects of adenine nucleotides fall within physiologically relevant concentrations, suggesting they may play a role in regulating the interaction of γ3-NTE with the α2-KD, although it remains unclear whether this effect is driven by γ nucleotide sites or the α-KD active site.

To further interrogate the γ3-NTE/α2-KD interaction we performed single cycle kinetics by surface plasmon resonance (SPR) using heterotrimeric *α*2β2*γ*1 AMPK or *α*2-KD(T172D) immobilised on a Ni^2+^-NTA sensor chip. Our SPR data revealed that the γ3-NTE interacted strongly with both active (CAMKK2-treated) and naïve (untreated) *α*2β2*γ*1 complexes, as well as with the isolated α2-KD (Fig. 3D-F). Kinetic analysis of the SPR sensorgrams, performed using the Biacore T200 Evaluation Software, showed that while the naïve complex exhibited the highest association (*k*_a_) and dissociation (*k*_d_) rates, the activated complex displayed the lowest *k*_d_ values. These kinetic parameters characterise the velocity of γ3-NTE binding to the complexes and stability of the resultant interaction, respectively. Consequently, the γ3-NTE demonstrated its highest affinity (lowest *K_D_*) for the active *α*2β2*γ*1 (Fig. 3G). These data imply that the role of the γ3-NTE is to switch off already activated AMPK, instead of maintaining naïve AMPK in an inactive state.

We next performed crosslinking experiments to confirm the involvement of the γ3 region between T130 and P168 in facilitating the γ3-NTE/α2-KD interaction. In these experiments, the zero-length crosslinker DMTMM (4-(4,6-dimethoxy-1,3,5-triazin-2-yl)-4-methylmorpholinium chloride) or photo-activatable crosslinker sulfo-SDA (sulfosuccinimidyl 4,4′-azipentanoate) (57) was incubated with purified γ3-NTE (AVI-tagged) and α2-KD. These reagents facilitate the formation of covalent bonds between residues in close spatial proximity, capturing both intra- and inter-protein interactions. Results required careful interpretation due to the similar molecular masses of the two proteins and propensity of both agents to form crosslinked α2-KD pairs that co-migrated with crosslinked γ3-NTE/α2-KD by SDS-PAGE (Fig. 4A, lane 4 & Fig 4B, lane 6). DMTMM-treated γ3-NTE also displayed slightly faster electrophoretic mobility than untreated γ3-NTE, presumably due to intra-protein crosslinks (Fig. 4A, lanes 1 and 2). Despite this, co-incubation of α2-KD and γ3-NTE with either agent resulted in clear increases in the abundance of a single crosslinked protein species relative to α2-KD alone (Fig 4A, lane 6 & Fig 4B, lane 7), with incorporation of both α2-KD and γ3-NTE in the DMTMM-crosslinked species confirmed by immunoblotting (Fig. 4C, lane 8). A γ3-NTE construct without an AVI-tag and the N-terminally shorter γ3_130-168-AVI_ fragment also successfully crosslinked with the α2-KD, confirming the interaction is specifically mediated via the γ3-NTE between T130 and P168 (Fig. 4C, lanes 4 and 10, & S6). Reduced intensity crosslinking bands were observed between α2-KD and the shortened C-terminal γ3_2-129_ fragment (Figs. 4C, lane 6, & S7), confirming the importance of the γ3_130-168_ region for the γ3-NTE/α2-KD interaction. The presence or absence of the MBP tag used during purification of the γ3-NTE constructs had no bearing on the ability of the γ3_130-168_ fragment to crosslink with α2-KD (Fig. S7). The greater molecular weight of the MBP tagged construct also provides physical separation from the minor species of α2-KD/α2-KD crosslinks present in these experiments, thereby confirming the presence of the γ3-NTE/α2-KD interaction (Fig. S7A-H). Our mass spectrometric analysis of the crosslinks yielded sparse data, with only a negligible number of unique crosslinks identified between the kinase domain and the γ3 NTE. Relying on these isolated observations would introduce significant selection bias. This structural interface remains technically challenging to capture via MS, necessitating future optimisation of our experimental conditions.

**Figure 4.**
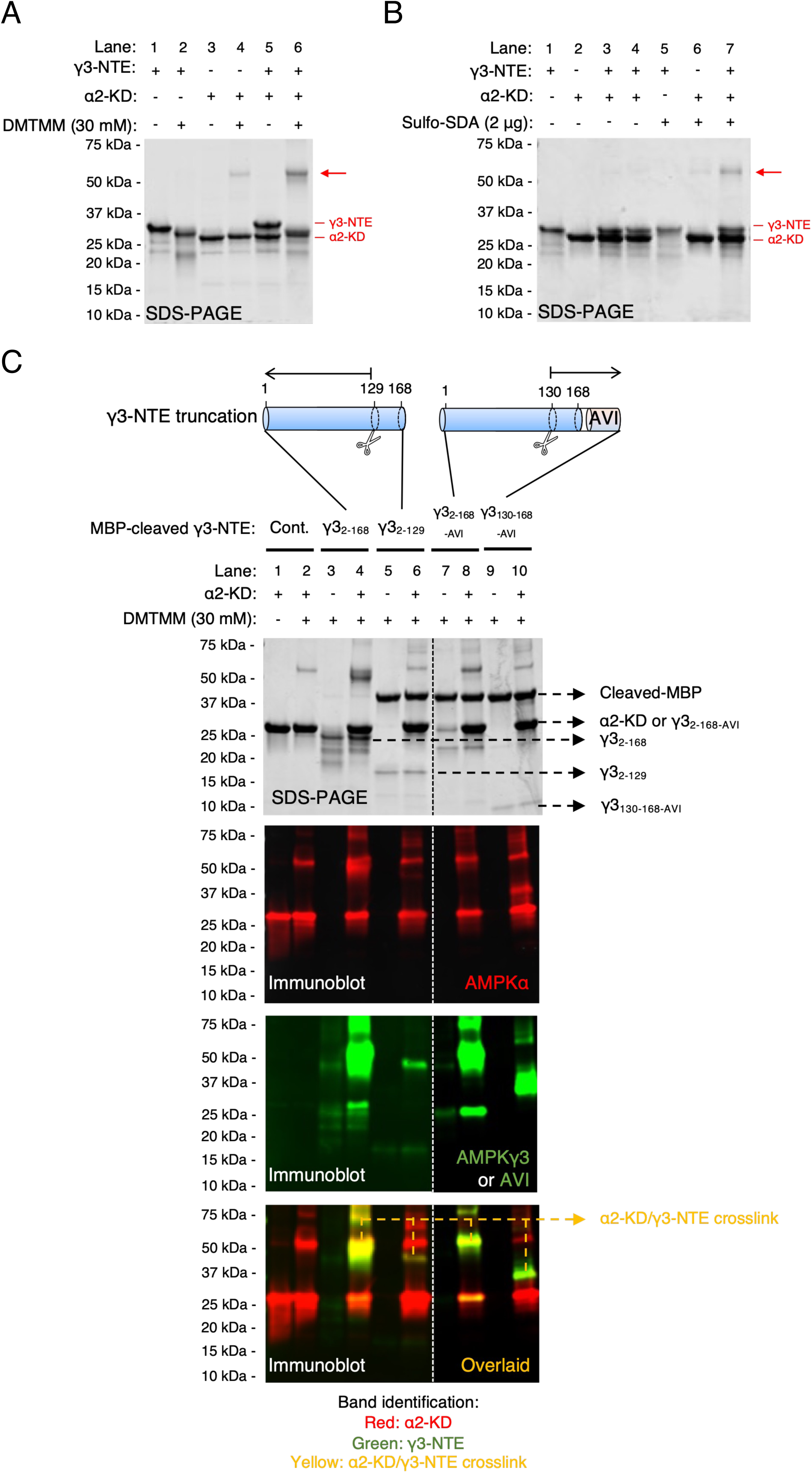
Chemical crosslinking of AMPK γ3-NTE with α2-KD. SDS-PAGE images of crosslinking reactions using (A) DMTMM and (B) sulfo-SDA. Red arrows indicate α2-KD/γ3-NTE crosslinked species. α2-KD and γ3-NTE bands are also indicated. (C) SDS-PAGE and immunoblot images of DMTMM-mediated crosslinking reactions between α2-KD and various MBP-cleaved γ3-NTE variants with different truncations. Red bands = α2-KD (AMPKα antibody), green bands = γ3-NTE (AVI-tag or AMPKγ3 antibody); yellow (merged) bands = α2-KD/γ3-NTE crosslinked species.

### The γ3-NTE interacts with the αC-helix of the α2-KD

We used hydrogen-deuterium exchange mass spectrometry (HDX-MS) to locate the site on the α2-KD that interacts with γ3-NTE (Figs. 5A & S8A). Incubation of α2-KD with γ3-NTE in a 1:1 molar ratio significantly reduced deuterium uptake for up to 1 minute on multiple peptides surrounding α2-KD residues 55-70 (^55^LDVVGKIKREIQNLKL^70^) (Fig. 5B-E). This protected region corresponds to the αC-helix (Fig. 5B), a major regulatory element for protein kinases (58) (Fig. 5F). There was no significant difference in deuterium uptake in any γ3-NTE peptides incubated with α2-KD compared to the control γ3-NTE in isolation (Fig. S8B), which we suspect is due to the lack of significant secondary structure that causes γ3-NTE peptides to become quickly saturated by deuterium at the initial time point of 6 seconds (Fig. S8C-H).

**Figure 5.**
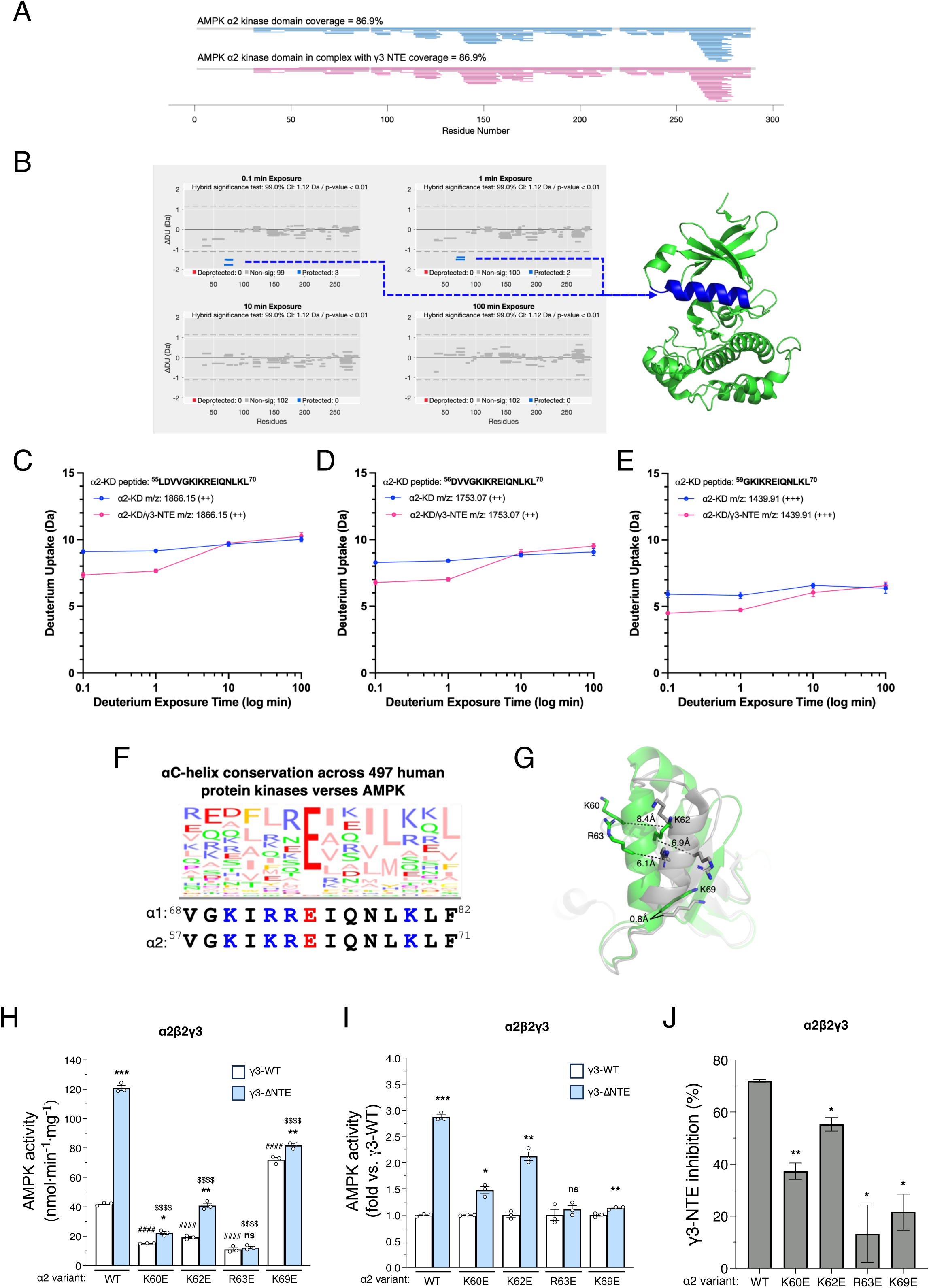
HDX-MS and mutational analysis reveal an interaction between AMPK γ3-NTE and α2-KD αC-helix. (A) Visualisation of HDX-MS protein sequence coverage for α2-KD. (B) Woods plots for differences in deuterium uptake (ΔDU) by α2-KD ± γ3-NTE across the three time points of deuterium exposure; 0.1 min, 1 min, 10 min and 100 min. A hybrid significance test consisting of a two-prong statistical test, implemented in Deuteros 2.0 was used to identify peptides that show a significant difference. Dashed horizontal lines indicate 99% confidence limit applied to the dataset to identify peptides with significant deuteration differences. Blue = significantly protected peptides (note upper band consists of 2 overlapping peptides); grey = unchanged peptides. *Right:* Differential HDX-MS data projected onto the crystal structure of AMPK α2-KD (green; PDB: 6B2E). Blue region represents deuteration-protected αC-helix (^56^DVVGKIKREIQNLKL^70^). Image was made in PyMOL. (C-E) Plots of deuterium uptake vs. exposure time for 3 deuteration-protected αC-helix peptides. Data shown as intensity-weighted mean values ± standard deviations (n=3). (F) Protein sequence of the αC-helices of α1 and α2 aligned with αC-helix conservation across 497 human protein kinases (82) displayed as a sequence logo generated in Jalview (83). (G) Cartoon representation of the active α2-KD N-lobe (green, PDB: 6B1U) superimposed on α1-KD in the inactive state (grey, PDB: 7JHG). Positively charged residues in the αC-helix are shown in stick representation. The inter-residue distances of K60, K63, R63 and K69 are measured from Cα-Cα. (H) Basal activities of AMPK α2β2-complexes (α2-KD WT and mutants) containing γ3-WT or γ3-ΔNTE. Data are shown as mean AMPK activity (nmol·min^−1^·mg^−1^) ± s.e.m.; *n* = 3. Statistical analyses were performed by one-way ANOVA with Dunnett’s multiple-comparisons test comparing to α2-WT complexed with γ3-WT, *^####^p* < 0.0001, or γ3-ΔNTE, *^$$$$^p* < 0.0001. (I) Basal activities of γ3-ΔNTE complexes in (H), expressed as fold-change relative to their respective γ3-WT complexes. Data are shown as mean fold-change in AMPK activity ± s.e.m.; *n* = 3. (J) Percentage inhibition exerted by the γ3-NTE calculated from (H). Data are expressed as the mean ± s.e.m.; *n* = 3. Statistical analyses were performed by Welch’s *t*-test vs. α2-WT complex. \**p* < 0.05, \*\**p* < 0.01. For (H & I), Statistical analyses were performed by Welch’s *t*-test vs. respective γ3-WT complexes. ns, *p* > 0.05, \**p* < 0.05, \*\**p* < 0.01, \*\*\**p* < 0.001.

The α2-KD αC-helix contains four positively charged basic residues; K60, K62, R63 and K69 (Fig. 5F), with crystal structures showing that K69 remains highly solvent-exposed and undergoes minimal structural deviation (∼0.8 Å) during the α-KD transition between active and inactive states (Fig. 5G). This conformational stability suggests that K69 may serve as a consistent docking point for the γ3-NTE. The γ3_130-159_ segment contains ten acidic residues (Fig. 6F) that could theoretically interact with basic αC-helix residues (Fig. 5F). To identify key residues involved in the γ3-NTE/α2-KD interaction, we individually mutated α2 αC-helix basic residues to acidic Glu and co-expressed them in HEK293T cells with β2 and either γ3-WT or γ3-ΔNTE (Figs. 5H and S9A). As previously observed (Fig. 1), deletion of the γ3-NTE elevated activity of WT α2 by 2.9-fold compared to γ3-WT, an effect that was reduced with α2 mutants K60E (1.5-fold), K62E (2.1-fold) and further reduced with R63E and K69E (only a 1.1-fold increase) (Fig. 5H, I, J). α2 mutations induced opposing effects on basal pT172 levels and activities of γ3-WT complexes. Specifically, compared to α2-WT, pT172 and activity was reduced 50-60% with the α2-K60E, α2-K62E and α2-R63E mutations, but increased 1.9-fold with the α2-K69E mutation (Figs. 5H & S7B). Similar to α2 WT, deletion of the γ3-NTE had no effect on pT172 in any of the α2 mutant complexes. Collectively, these data indicate a potential γ3-NTE docking site on the α2 αC-helix and provide insights on its inhibitory mechanism. Of note, charge swapping at α2 position 69 increased basal activity that was not further increased with γ3-NTE deletion, indicating this residue is a possible key determinant in the γ3-NTE/α2-KD interaction (Fig. 5G).

**Figure 6.**
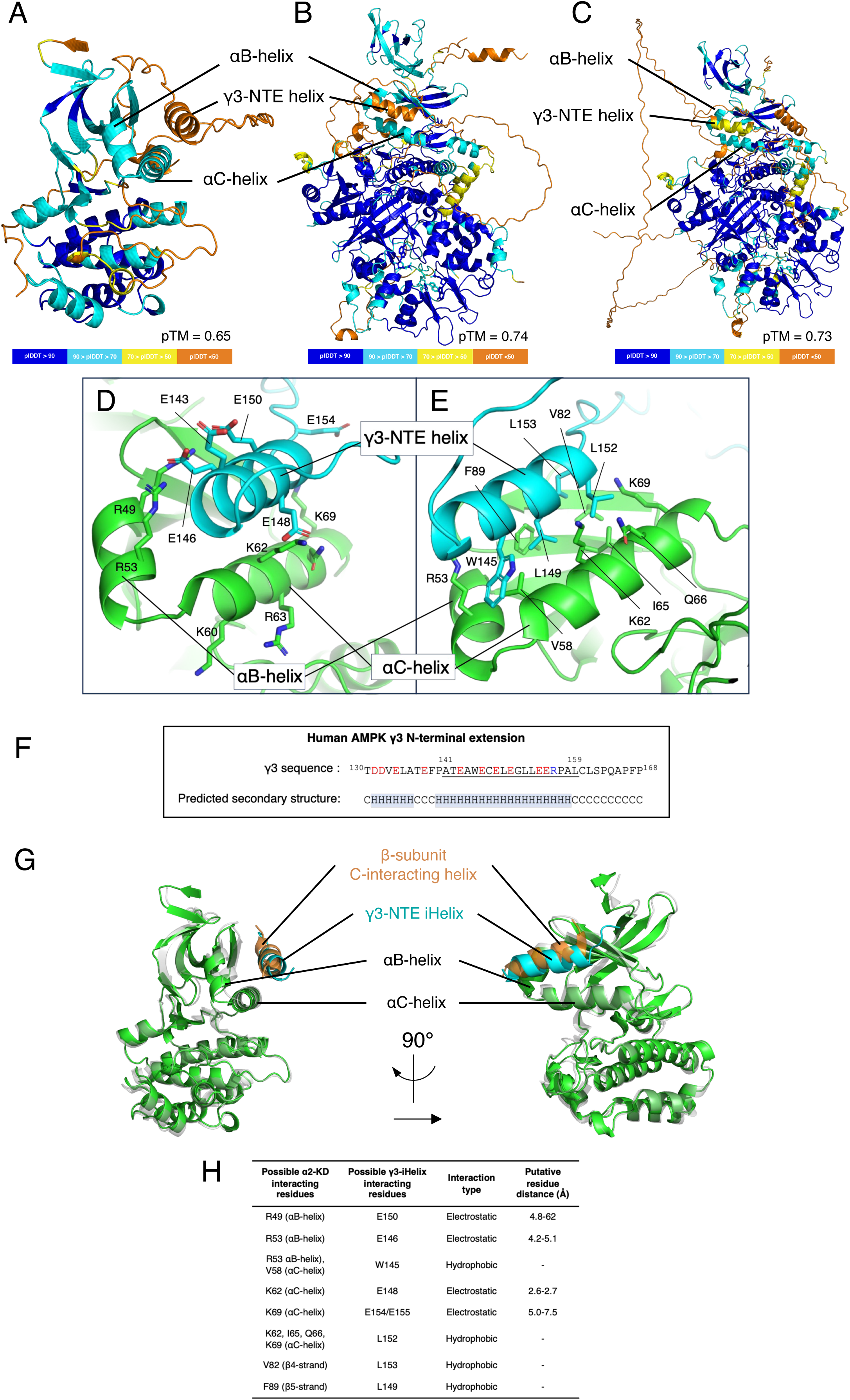
AlphaFold3 visualisation of the interaction between AMPK γ3-NTE and α2-KD αC-helix. Cartoon representation of AlphaFold3-predicted AMPK structures with predicted local distance difference test (plDDT) score per residue colour projected onto the structure according to the colour confidence indicator shown: blue, very high; cyan, high; yellow, low; and orange, very low. (A) Predicted γ3_95-178_ interaction with α2_6-278_, predicted template modelling (pTM) score = 0.65. (B) Predicted γ3_95-178_ interaction with α2β2γ3_179-489_, β2 residues 159-171 were mutated to glycine residues, pTM score of 0.74. (C) Predicted full-length trimeric AMPK α2β2γ3, with β2 residues 159-171 and γ3 residues 172-177 mutated to glycine, pTM score of 0.73. Close up view of (A) showing (D) charged residues and (E) hydrophobic residues of the αB/αC- and γ3-NTE helices, forming a complimentary binding surface. (F) Protein sequence of γ3 residues 130-168 with predicted secondary structure according to the Chou and Fasman secondary structure prediction algorithm (61). Red = acidic residues; blue = basic residues. (G) Superimposed AlphaFold3 structure (green and cyan) from (A) with the crystal structure of α2-KD (gray) and β-subunit C-interacting helix (orange) (PDB: 6B2E). (H) List of possible key interacting residues between α2-KD and γ3-NTE with their respective predicted molecular distances derived from models in (A-C).

### Structural predictions of the γ3-NTE binding to the α2-KD

As experimental structural data for the γ3-NTE remains unavailable via cryo-EM or crystallography, we employed AlphaFold3 (via the AlphaFold Server) to predict the γ3-NTE binding to the α2-KD and visualise the potential structural determinants driving this interaction (59, 60). The confidence level for each predicted model was determined from the integrated AlphaFold confidence metrics. The first is the predicted local distance different test (pLDDT), which calculates values ranging from 0 to 100 for each residue, with 100 the highest confidence. The pLDDT confidence range is shown as colour outputs on the image structure. The second metric is predicted template modelling (pTM) that ranges from 0 to 1. pTM is a confidence metric measuring the similarity between the modelled complex and the hypothetical “true” structure. Values above 0.5 predicts the complex is likely to be indicative of a biologically relevant structure. The top-ranked models assessed by the pLDDT and pTM were chosen for viewing on PyMOL unless otherwise stated (Fig. 6A, B, C).

Initially we modelled α2_6-278_ and γ3_2-178_ as a complex, however this resulted in an unstructured γ3-NTE (data omitted for clarity). Some of the known phosphorylation sites on the γ3-NTE, such as γ3-S14 and γ3-S65 (46), were placed in the α2-KD substrate-binding groove which may have physically prevented the γ3_130-159_ segment from reaching the αC-helix. In support of this, replacing γ3_2-178_ with a shorter γ3_95-178_ sequence (Table S1) revealed a helical structure on residues A141-L152 positioned within sidechain-interacting distance of the αC-helix (Fig. 6A, D, E; pTM = 0.65). In support of this modelling, we used the Chou and Fasman secondary structure prediction algorithm (61), which also predicted a helical structure in this region of the γ3-NTE on residues P140-A158 (59, 61) (Fig. 6F). Our biochemical data has identified an inhibitory motif that overlaps with the predicted helical region, and as such we have termed this γ3 helix the γ3-inhibitory-helix (γ3-iHelix). The predicted interaction model of the α2-KD with the γ3-iHelix is akin to that of the experimentally determined C-interacting helix of the β-subunit in ligand-bound, active AMPK crystal structures (Fig. 6G); for example, in the 6B2E PDB model (36).

Next, we modelled the heterotrimeric α2β2γ3_169-489_ in complex with the truncated γ3-NTE, γ3_95-168_ (Table S1). Initially, the γ3_95-168_ sequence was unstructured and not bound to the αC-helix (data not shown), likely due to high confidence positioning of the β-subunit C-interacting helix bound to the αC-helix in an orientation closely resembling experimentally determined AMPK crystal structures that are used for AlphaFold3 training. To circumvent this, we replaced the C-interacting helix sequence with a flexible glycine linker, (β2_D159-M171_→G). Using this linker, AlphaFold3 predicted a helical region corresponding to the γ3-iHelix (forming on residues T142-E154) bound to the αC-helix in place of the C-interacting helix (Fig. 6B; pTM = 0.74), closely resembling the α2_6-278_/γ3_95-178_ structure outlined above (Fig. 6A, D).

As we used isolated γ3 sequences for the aforementioned structural predictions, we wanted to determine whether the γ3-iHelix (∼T142-E154) could physically interact with the αC-helix within a heterotrimer AMPK complex, located approximately 50Å (measured linearly) from the end of the γ3-NTE sequence (γ3-I183). The ∼30-amino-acid sequence between the γ3-iHelix and the CBS domains could theoretically traverse ∼105 Å if fully extended into linear chain, which is more than sufficient to bridge the 50 Å gap. However, in a compact helical conformation, this same region would span only ∼45 Å, rendering the αC-helix structurally inaccessible. Therefore, whilst it is plausible the γ3-iHelix could reach the αC-helix of the α-KD, there are some structural elements situated between the γ3 pre-CBS1 sequence and the αC-helix, most notably the AID, which could sterically obstruct its path. To test this from a theoretical standpoint, we aimed to visualise the predicted full-length α2β2γ3 AlphaFold3 structure (Table S1). Initially, we modelled heterotrimeric α2β2γ3 with a mutated C-interacting helix (β2_D159-M171_→G), with the resulting γ3-NTE predicted to be mostly unstructured (data not shown). However, we noticed in this structure that γ3 residues G171-R177, immediately preceding the γ3 pre-CBS1 sequence, formed a short helix directing the γ3-NTE away from α2-KD (data not shown). To examine the consequences of removing this helix, we re-modelled heterotrimeric α2β2γ3 as above with the addition of a small flexible glycine linker (γ3_W172-R177G_→G), the resulting prediction placing the γ3-iHelix (γ3_E143-E154_) next to the αC-helix (Fig. 6C; pTM = 0.73) in line with our previous models (Fig. 6A, B). This raises the question of whether this short helix that precedes the γ3 pre-CBS1 sequence plays a regulatory role for the γ3-iHelix.

Deeper analysis of our predicted structures revealed several positively charged α2-KD residues (R49, R53, K62, and K69) situated on the on αB- and αC-helices interacting with the negatively charged γ3-iHelix residues (E146, E148, E150, E154 and E155) (Fig. 6D, H). Some hydrophobic α2-KD residues (V58 and I65) are also positioned to face and interact with hydrophobic residues of the γ3-iHelix (W145, L149 and L152, Fig. 6E, H). Overall, the modelling provides a plausible interaction surface for γ3-iHelix binding to the α2-KD via residues on the αB- and αC-helices. While we have not elucidated the complete inhibitory mechanism, we identified residues that likely play an important role in γ3-iHelix binding to the α2-KD to inhibit AMPK activity.

### The γ3-NTE contains at least three phosphorylation sites

All subunit isoforms of AMPK are subject to phosphorylation on distinct sites, many of which with have known regulatory effects (37), although very little is known regarding how phosphorylation affects γ3-AMPK activity. Therefore, we sought to identify novel phosphorylation sites that could be involved in the regulation of the γ3-iHelix. Here, we applied LC-MS/MS to tryptic digests of HEK293T cell-expressed α2β2γ3 and confirmed previous reports that S14 and S65 are γ3 phosphorylation sites (Fig. S10A, B) (46, 47), with their signals abolished by the exchange of each site for a non-phosphorylatable Ala (Fig. S10D, E). As previously reported, we found that the γ3_96-169_ peptide was too large for tryptic LC-MS/MS analysis (46), likely due to blocked cleavage by Arg-Pro at position 156-157. Despite using alternative proteolytic strategies, we were unable to achieve reliable coverage of this sequence. To circumvent this issue, we mutated γ3-D131 to Lys which allowed us to detect two peptides covering the γ3_96-169_ region. We were unable to detect any phosphorylation sites within the γ3_96-131_ peptide (data not shown); however, appreciable level of phosphorylation of the γ3_132-169_ peptide were detected, with MS/MS data and subsequent substitution of S162 with an Ala indicating γ3-S162 was indeed the phosphorylated residue (Fig. S10C, F). Stoichiometry measurements using area under the curve analysis of the peptide peaks (without normalisation) indicated that the basal phosphorylation of γ3-S162 is ∼16%, which approximates previously measured stoichiometries of several functional phosphorylation sites on AMPK (46). Of the three detected phosphorylation sites situated on the γ3-NTE, γ3-pS162 is the most proximal to the γ3-iHelix (∼T142-E154), making it a strong candidate for a ‘phospho-switch’ toggling γ3-iHelix-mediated AMPK activity. The proline at the P+1 position serves as a docking motif for a broad range of (proline-directed) protein kinases, many of which are implicated in metabolic stress responses provoked by physiological stimuli like exercise.

## Discussion

The central finding of this study is the identification of an autoinhibitory sequence encoded in the NTE of the γ3 subunit isoform of AMPK that we have termed the γ3-iHelix (residues T142-E154). This helix interacts with the αC-helix of the kinase domain of AMPK, which may play a role in maintaining the low basal activity levels of predominantly muscle-expressed α2β2γ3 complexes, independent of changes in activation loop α-T172 phosphorylation. Our *in cellulo* phosphoproteomic profiling confirms that the loss of the γ3-NTE shifts the cellular kinase toward an active state marked by the increase in β2-S108 autophosphorylation (37). However, the activity shift caused no change to the autophosphorylation of the inhibitory ST-loop site α2-S491 (37). In fact, α2β1γ3 complexes exhibited a distinct reduction in S491 phosphorylation. This implies γ3-ΔNTE enhances AMPK catalytic output, while simultaneously reducing the accessibility of the ST-loop to its own kinase domain, preventing the typical self-limiting autophosphorylation mechanism. Furthermore, the widespread reduction in phosphorylation on known mTORC1 (α1/2-S347/345, α2-S377) and GSK3β (α2-S481, α1/2-T481/485) inhibitory sites provides strong evidence of a cellular feedback mechanism. Elevated AMPK activity likely triggers the well-characterised phosphorylation of RAPTOR and TSC2, suppressing upstream mTORC1 signalling (62, 63) and potentially downregulating GSK3β activity (53). By systematically dampening these upstream inhibitory inputs, the cell reinforces and sustains the increased activity state of the γ3-ΔNTE complex.

The γ3-iHelix adds a secondary layer of regulation on top of the α-subunit AID, a tri-helical bundle that binds the hinge region of the kinase domain between the αC and αE-helices to allosterically inhibit AMPK (64), but dissociates upon AMP/ADP binding to the γ-subunit. The existence of the γ3-iHelix would suggest there is a need to discretely regulate the activity of γ3-containing AMPK complexes independently of cellular adenine nucleotide ratios (AMP/ADP:ATP). One possibility is that the γ3-iHelix allows γ3-containing AMPK complexes to regulate their own activity in response to metabolic stresses (e.g., glycogen deprivation, oxidative stress) generated in skeletal muscle by strenuous physical exercise (37). γ3-AMPK complexes in human skeletal muscle are by far the most sensitive to increased contractile activity, especially high-intensity modalities (17, 65). In this context, γ3-AMPK is essential for insulin-independent glucose uptake immediately following exercise, and γ3-AMPK activity can remain elevated for up to three hours following the exercise bout (41, 66, 67). Since adenine nucleotide concentrations return to basal levels within minutes after cessation of exercise, it follows that the regulation of γ3-AMPK extends beyond that of canonical nucleotide sensing. There is evidence that AMP has a higher affinity for CBS site 1 in γ3 than in γ1 (48), although it is widely accepted that γ3 complexes are largely resistant to allosteric activation by AMP (45). AMP may remain bound to γ3 to maintain AMPK activity through preservation of pT172 (41), whereby tuneable γ3-AMPK activity is conferred by the γ3-iHelix. Tuneable activation of γ3-AMPK during exercise may be the result of relieving γ3-iHelix autoinhibition, perhaps via a post-translational modification(s) like phosphorylation, or another reversible modification. Whether the γ3-iHelix endows γ3-AMPK complexes with the intrinsic ability to self-regulate independent of disruptions to the adenine nucleotide pool remains a matter for future work.

The persistence of elevated AMPK activity by truncation of the γ3-NTE (γ3-ΔNTE) indicated that its increased activity was not completely dependent on its increased affinity for ATP (*K*_M_). Although mammalian γ3-ΔNTE AMPK complexes display mildly reduced *K*_M_ for ATP compared to the γ3-WT, bacterial γ3 complexes have a significant 10-fold increase in the *K*_M_ for ATP compared to γ1 complexes, in line with previous reports (48). This discrepancy between expression systems likely underscores the regulatory impact of mammalian-specific post-translational modifications or β-myristoylation, both of which are absent in bacterial preparations but critical for modulating AMPK activity (37).

β1 and β2 isoforms possess distinct sequence differences within their CBM and linker regions, with β1-subunit isoform-containing complexes generally more responsive to ADaM-site activators, despite both β-isoforms utilising an identical kinase activating mechanism (34, 35, 50, 68–71). Our kinetic data comparing the impact of α1/2 and β1/2 isoforms on γ3-AMPK complex activity ± γ3-NTE, reveals an interesting phenomenon; α1β2 and α2β1 complexes are more susceptible to γ3-iHelix inhibition, as its deletion triggers a ∼7.5-fold increase in the catalytic efficiency relative to a ∼3.5-fold for α1β1 and α2β2 complexes. This suggests α/β-subunit interactions play a role in the inhibitory strength of the γ3-iHelix, likely due to small conformational changes between the different α/β isoform combinations. Our proposed site of interaction for the γ3-iHelix is the αC-helix, which structurally overlaps with the β-subunit C-interacting helix binding site, visualised in active, ADaM ligand-bound crystal structures of AMPK (72). Therefore, we propose an autoinhibitory mechanism whereby the γ3-iHelix binds to the αC-helix, concurrently hindering AMPK activation by β-subunit C-interacting helix and promoting a structural re-arrangement (outward rotation) of the αC-helix that disrupts the position of key catalytic residues. This proposal is strongly supported by the introduction of individual charge-repulsion mutations to the four conserved basic residues of the αC-helix, with the predominant effect elicited by α2-K69E. The four mutations reduced the γ3-NTE inhibition in the α2β2γ3 complex. We acknowledge that the introduction of negatively charged acidic residues within the basic αC-helix is “aggressive” with all but α2-K69E displaying impaired kinase function compared to their WT γ3-AMPK counterparts. Importantly, the elevated specific activity of α2-K69E mutation in γ3-AMPK is in line with γ3-ΔNTE, suggesting that the inhibitory effects of the γ3-NTE have been reduced by this single point mutation. We propose the increase in basal activity of the mutant is due to the release of the γ3-NTE inhibitory mechanism by the introduction of the negatively charged glutamic acid residue on the outer surface of the αC-helix, thereby repelling the highly acidic γ3-iHelix from forming electrostatic interactions with the αC-helix. While the basic αC-helix residues may have individual roles in coordinating binding of the γ3-NTE, it remains unclear if all four residues are required for direct binding of the γ3-NTE.

The AlphaFold3 models corroborate our extensive *in vitro* biochemical data, firstly by predicting the helical conformation of the γ3-iHelix and second, by revealing the potential for α-subunit αC-helix residues to act as docking sites for the negatively charged γ3-iHelix. Structural analysis of the αC-helix in inactivated AMPK reveals the α2-K69E residue is solvent-exposed and thus could serve as a docking site for the γ3-iHelix. However, in structures of activated AMPK, access to the αC-helix is partially occluded by the β-subunit C-interacting helix. Our SPR analysis shows that the γ3-NTE has a higher affinity for the AMPK heterotrimer than the isolated α2-KD, particularly the active, τ172-phosphorylated AMPK complex, indicating the γ3-iHelix can outcompete the β-subunit C-interacting helix to interact with the αC-helix to keep AMPK activity in check. Importantly, α2-K69 is located at the end of the αC-helix and its charged side chain remain accessible even in the presence of a bound β-subunit C-interacting helix. The lower affinity for monomeric α2-KD suggests the γ3-NTE has additional binding sites outside of the kinase domain. This observation is supported by our initial γ3-NTE truncation experiment (Fig. 1E), which revealed a small but significant increase in activity when the truncations increased from residue W60 to P95. A prior study demonstrated that in response to prolonged (24 h) glucose starvation in mammalian NIH-3T3 cells, the τ172-phosphorylated α-subunit was more susceptible to proteasomal degradation than non-phosphorylated species, presumably to safeguard against AMPK-mediated, unrestrained autophagic flux that can be injurious (73). Hence, the affinity γ3-iHelix has for active AMPK complexes is likely indicative of some form of homeostatic fine-tuning to preserve metabolic balance following exposure to bioenergetic stress (i.e., that which is caused by exercise).

The αC-helix is an allosteric regulatory hotspot for protein kinases (30). In active AMPK, the αC-helix is structurally supported by interactions with pT172 in the activation loop (19, 42), maintaining αC-helix residue α2-K69 solvent exposed and accessible for γ3-iHelix interaction. Our SPR data indicates the γ3-iHelix can displace the β-subunit C-interacting helix, forming a new γ3-iHelix/αC-helix interaction, potentially rotating the αC-helix outward to emulate the inactive ATP-bound cryoEM structure of AMPK (PDB ID: 7JHG) (74). In the inactive conformation of AMPK, the αC-helix is rotated, and several key catalytic structures/interactions are disrupted, such as the regulatory spine that controls the ON/OFF-switch in protein kinases (42, 74). To formally validate this hypothesis, a high-resolution structure of α2β2γ3 capturing the γ3-iHelix bound αC-helix is required.

We assessed the phosphorylation status of the γ3-NTE under basal cellular conditions and in addition to the previously identified γ3-pS14 and γ3-pS65 sites, identified a novel phosphosite, γ3-pS162 (46). We predict γ3-S162 could be a regulatory site given its structural proximity to the γ3-iHelix. The sequence surrounding γ3-S162 (RPALCL**<u>S</u>**^162^PQAPFP) is noteworthy being a Ser-Pro site, as these are abundant and consistently phosphorylated across most AMPK subunits, in particular the γ2 isoform (46). All three γ3 phospho-sites likely have important functional roles. For instance, a recent study found that γ3-pS65 was strongly associated with insulin resistance in human skeletal muscle, yet its dephosphorylation did not affect basal, MK8722- or H_2_O_2_-induced AMPK activity (47). It is plausible that γ3-NTE phosphorylation controls the ON/OFF-switch dynamics of AMPK. Whether this is a combinatorial effort involving the convergence of divergent signalling pathways, or alternatively driven by a single phospho-switch, is an area open for future research.

In summary, our study has revealed an inhibitory function for the AMPK γ3-NTE, which binds to the αC-helix of the α-KD to suppress AMPK activity. The significance of these findings is underscored by the fact that γ3-containing complexes are found almost exclusively in skeletal muscle and due to their prominent role in glucose homeostasis, makes them highly sought after targets for the treatment of T2DM. γ3-selective activators would bypass any deleterious effects seen in other tissues (i.e., cardiac hypertrophy) which is associated with systemic AMPK activation. As such, this new information will assist in the future development of isoform-specific AMPK therapeutics.

## Methods

### Plasmid constructs, mutagenesis and cloning

For mammalian cell expression, sequences for human wild-type γ3 and N-terminally truncated γ3 were generated with a C-terminal FLAG-tag and cloned into pcDNA3.1(−) using NotI/EcoRI restriction sites by Gene Universal (Newark, Delaware, United States). Other N-terminally truncated versions of the human γ3 construct were cloned into pcDNA3.1(−) using NotI/EcoRI restriction sites and primers outlined in Table 2 or by Gene Universal (Newark, Delaware, United States) as stated in Table 3. For bacterial expression, the human DNA sequence for the γ3 N-terminal fragments were generated with an N-terminal MBP-tag (PreScission cleavable) and C-terminal AVI-tag, if stated, and cloned into pMAL-c5X using NdeI/BamHI restriction sites by General Biosystems (Table 4). For heterotrimeric γ3 bacterial expression, the human DNA sequence for γ3 was cloned into pET DUET-1 multiple cloning site (MCS) 2 (MCS1 already containing _6xHis_α2) using MfeI/XhoI restriction sites, replacing the γ1 DNA sequence from our previous study (75) using the primers detailed in Table 2. It should be noted that the cloning strategy used to create this construct introduced additional N-terminal residues (M → MADLNW) on the γ-subunit, while the extreme C-terminus of γ3 is an alternate sequence (DALGA → DPSGPEKI) that was originally described by Cheung et al. (76). All other plasmids used in this study have been described previously (35, 70, 75, 77).

**Table 2.**
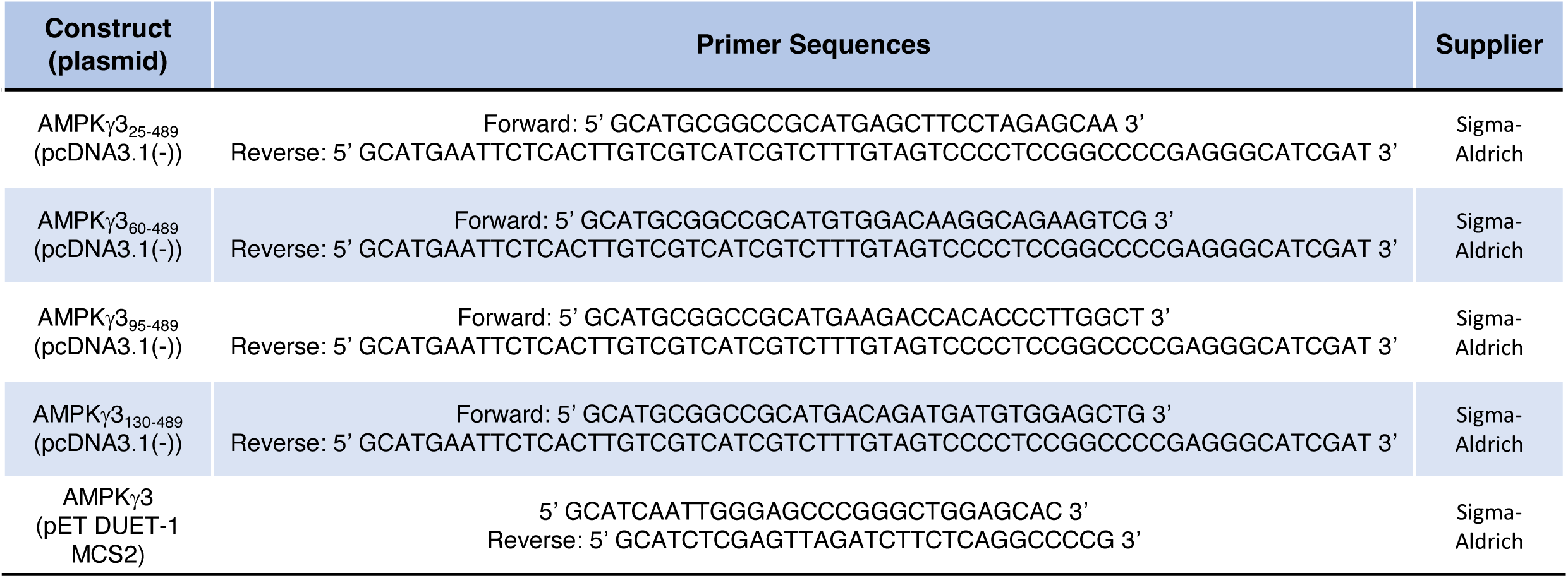
Details of single-stranded DNA primers used for cloning.

**Table 3.** AMPK plasmids in pcDNA3.1(−) vector used for expression and purification of recombinant AMPK complexes in mammalian cells.

| Construct | Note/Supplier |
| --- | --- |
| $\alpha 1$ | (77) |
| $\alpha 2$ | (77) |
| $\alpha 2^{\text{K60E}}$ | This study |
| $\alpha 2^{\text{K62E}}$ | This study |
| $\alpha 2^{\text{R63E}}$ | This study |
| $\alpha 2^{\text{K69E}}$ | This study |
| $\beta 1_{\text{MYC}}$ | (77) |
| $\beta 2_{\text{MYC}}$ | (77) |
| $\gamma 1_{\text{FLAG}}$ | This study |
| $\gamma 3_{\text{FLAG}}$ | Gene Universal |
| $\gamma 3_{26-489\text{-FLAG}}$ | Gene Universal |
| $\gamma 3_{60-489\text{-FLAG}}$ | Gene Universal |
| $\gamma 3_{95-489\text{-FLAG}}$ | Gene Universal |
| $\gamma 3_{110-489\text{-FLAG}}$ | Gene Universal |
| $\gamma 3_{120-489\text{-FLAG}}$ | Gene Universal |
| $\gamma 3_{130-489\text{-FLAG}}$ | Gene Universal |
| $\gamma 3_{140-489\text{-FLAG}}$ | Gene Universal |
| $\gamma 3_{150-489\text{-FLAG}}$ | Gene Universal |
| $\gamma 3_{160-489\text{-FLAG}}$ | Gene Universal |
| $\gamma 3_{169-489\text{-FLAG}}$ | Gene Universal |
| $\gamma 3_{179-489\text{-FLAG}}$ | Gene Universal |
| $\gamma 3_{185-489\text{-FLAG}}$ | Gene Universal |
| $\gamma 3_{\text{FLAG}}^{\text{S14A}}$ | This study |
| $\gamma 3_{\text{FLAG}}^{\text{S65A}}$ | This study |
| $\gamma 3_{\text{FLAG}}^{\text{D131K}}$ | Gene Universal |
| $\gamma 3_{\text{FLAG}}^{\text{D131K+S162A}}$ | This study |

**Table 4.**
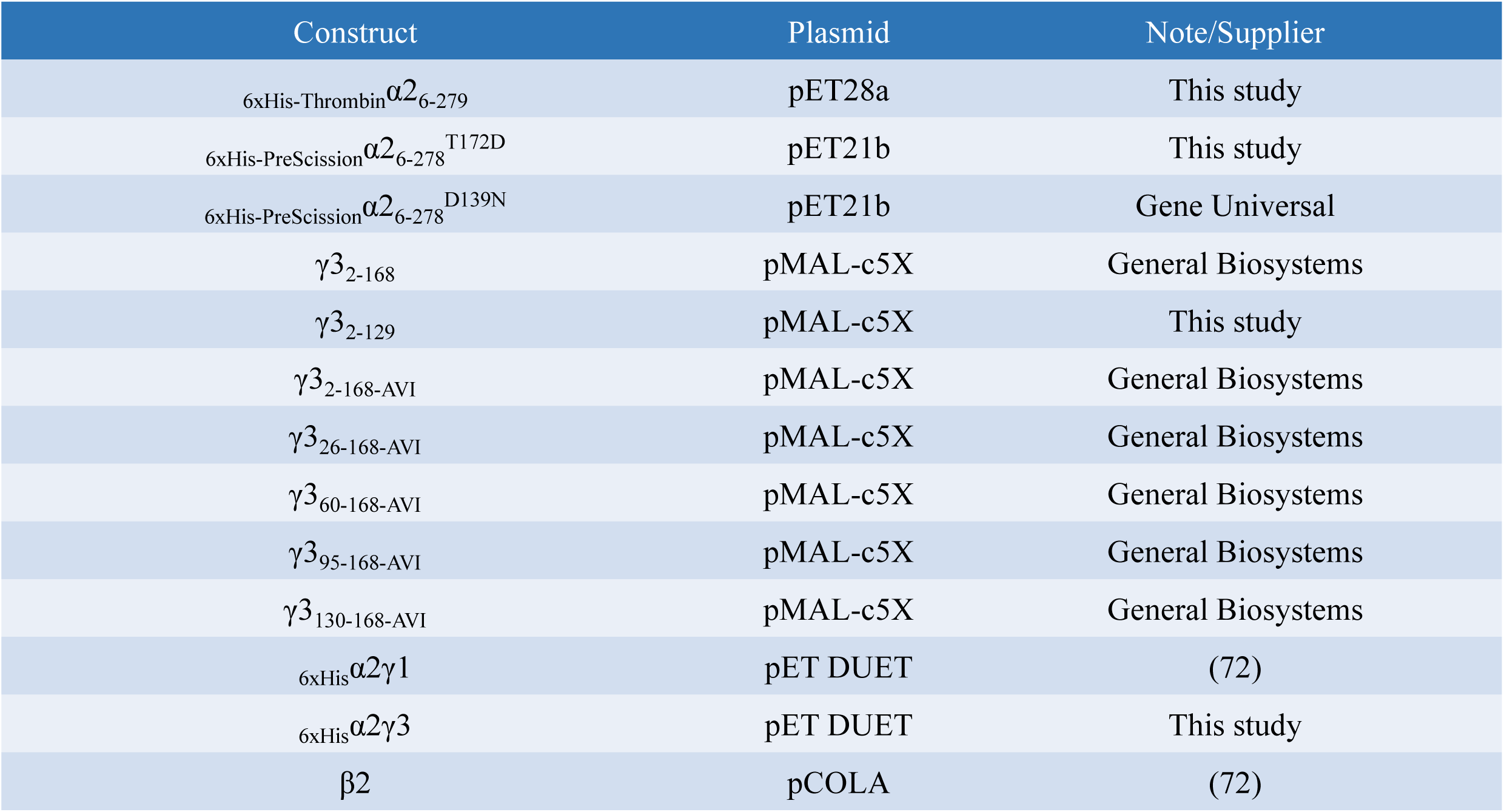
AMPK plasmids used for bacterial expression.

For mutagenesis, primers containing the desired mutation (detailed in Table 5) were designed and ordered from Sigma-Aldrich. Site-directed mutagenesis was carried out as previously described (37). Briefly, the DNA was amplified by PCR using a thermocycler, Dpn1 (NEB) was added to digest methylated parental DNA and the amplified DNA transformed into α-select competent cells (Bioline). Individual colonies grown on Luria-Bertani broth (LB) agar plates containing 100 µg·mL^−1^ antibiotics were inoculated in 5-10 mL LB containing 100 µg·mL^−1^ antibiotics and incubated overnight at 37 °C. Plasmid DNA was isolated using a Wizard Plus SV minipreps DNA purification kit (Promega) as per the manufacturers protocol. All new constructs were confirmed by Sanger sequencing (Australian Genome Research Facility, Melbourne, Australia).

**Table 5.**
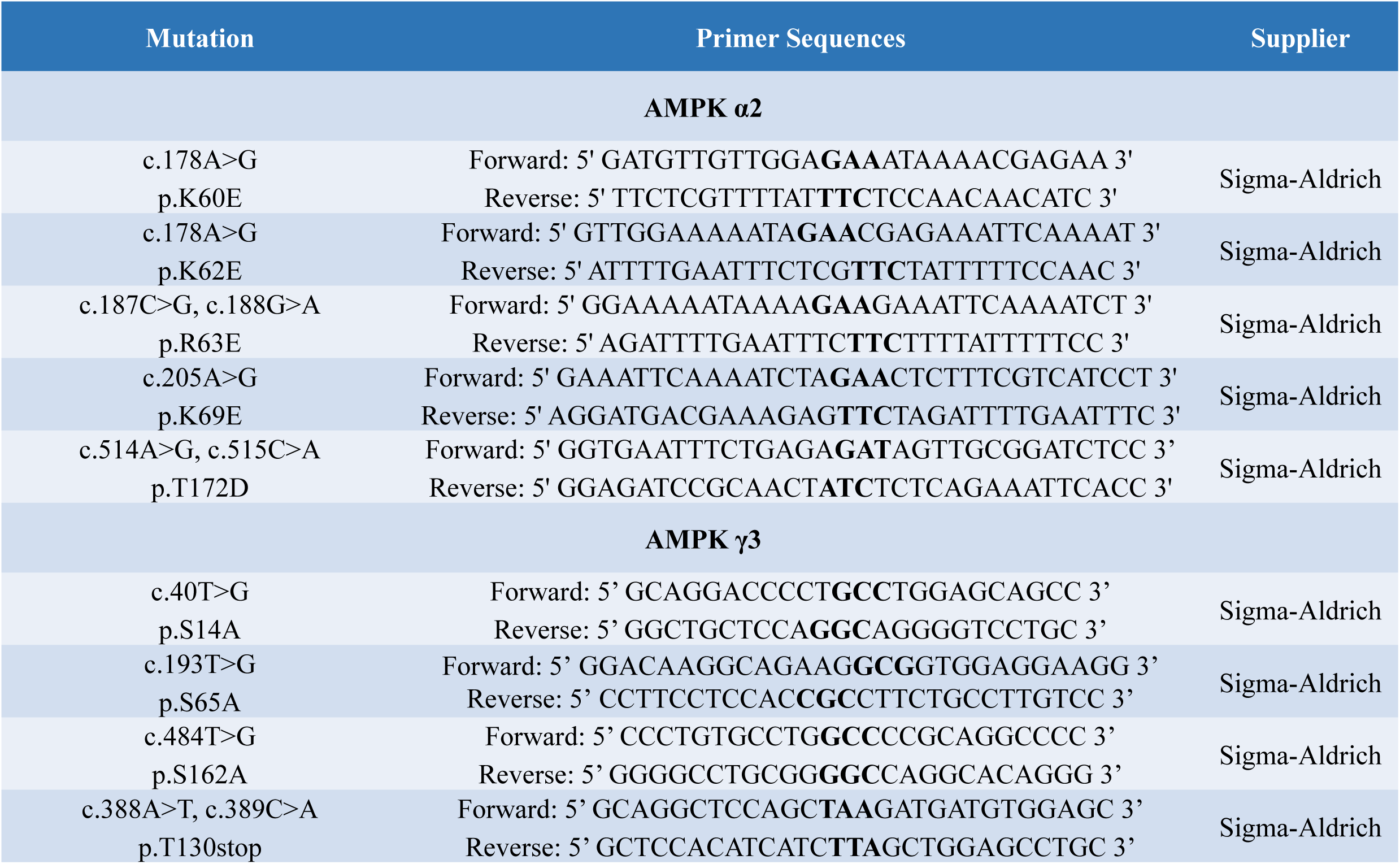
Details of single-stranded DNA primers used for site-directed mutagenesis. The mutating codons on the primer sequences are in bold.

### Immunoblotting and Coomassie gels

Protein samples were separated on gradient SDS-PAGE gels (Bio-Rad). For visualising total protein content, the gel was stained in Coomassie blue. For immunoblotting, the protein was transferred to an Immobilon-FL PVDF membrane (Merck Millipore). After blocking the membrane with 2% non-fat dry milk dissolved in PBS + 0.1% (v/v) Tween-20 (PBST; Sigma-Aldrich), membranes were incubated with primary antibodies for either 2 hours at room temperature or overnight at 4 °C, as indicated in Table 6. Following repeated washes with PBST, fluorescently labelled secondary antibodies diluted in PBST were added, if required, as detailed in Table 6. The membranes were then washed in PBST and visualised on the Odyssey Infrared Imaging System (LI-COR Biosciences) and immunoreactive bands analysed and quantified using Image Studio Software (LI-COR Biosciences).

**Table 6.** List of primary and secondary antibodies used for immunoblotting.

| Antibody<br>(Catalogue Number) | Type | Source | Incubation Period | Dilution | Supplier |
| --- | --- | --- | --- | --- | --- |
| AMPKα (2793) | Primary | Mouse | 2 hours/overnight | 1:1000 | Cell Signaling Technology |
| AMPKα1/2-pT174/T172 (2535) | Primary | Rabbit | 2 hours/overnight | 1:1000 | Cell Signaling Technology |
| AMPKα1/2-pS487/S491 (2185) | Primary | Rabbit | Overnight | 1:1000 | Cell Signaling Technology |
| AMPKβ (4150) | Primary | Rabbit | Overnight | 1:1000 | Cell Signaling Technology |
| AMPKγ3 (2550) | Primary | Rabbit | 2 hours/overnight | 1:1000 | Cell Signaling Technology |
| AVI Tag (A00674) | Primary | Rabbit | Overnight | 1:1000 | GenScript |
| FLAG Tag (9146) | Primary | Mouse | 2 hours/overnight | 1:1000 | Cell Signaling Technology |
| FLAG Tag (14793) | Primary | Rabbit | 2 hours/overnight | 1:1000 | Cell Signaling Technology |
| MYC Tag (2276) | Primary | Mouse | 2 hours/overnight | 1:1000 | Cell Signaling Technology |
| β-Actin (4970) | Primary | Rabbit | 2 hours | 1:1000 | Cell Signaling Technology |
| α-Tubulin (3873) | Primary | Mouse | 2 hours | 1:1000 | Cell Signaling Technology |
| ACC-pS79 (3661S) | Primary | Rabbit | Overnight | 1:1000 | Cell Signaling Technology |
| IRDye® 680 RD Streptavidin (926-68079) | Primary | - | Overnight | 1:1000 | LI-COR Biosciences |
| Alexa Fluor® 790 IgG Fraction Monoclonal Mouse Anti-Rabbit IgG, light chain specific (211-652-171) | Secondary | Mouse | 1 hour | 1:10,000 | Jackson ImmunoResearch Laboratories Inc. |
| Alexa Fluor® 680 AffiniPure Goat Anti-Mouse IgG, light chain specific (115-625-174) | Secondary | Goat | 1 hour | 1:10,000 | Jackson ImmunoResearch Laboratories Inc. |
| IRDye 680RD (926-68070) | Secondary | Mouse | 1 hour | 1:10,000 | LI-COR Biosciences |
| IRDye 680RD (926-68071) | Secondary | Rabbit | 1 hour | 1:10,000 | LI-COR Biosciences |
| IRDye 800CW (926-32210) | Secondary | Mouse | 1 hour | 1:10,000 | LI-COR Biosciences |
| IRDye 800CW (926-32211) | Secondary | Rabbit | 1 hour | 1:10,000 | LI-COR Biosciences |

### Cell culture

HEK293T/17, HEK293T, and COS7 cells were all maintained in Dulbecco’s Modified Eagle’s medium (DMEM; Sigma-Aldrich, D5796) supplemented with 10% fetal bovine serum (FBS; Assay Matrix) at 37 °C and 5% CO_2_.

### Mammalian protein expression and purification

Adherent HEK293T/17, HEK293T, or COS7 cells at ∼50% confluency were triply transfected with full-length α1 or α2, or α2 mutants (untagged), β1 or β2 (C-terminal MYC fusion), and full-length or truncated γ3 or γ3-mutants (C-terminal FLAG fusion) with FuGENE HD transfection reagent (Promega) as per the manufacturers protocol. Cells were harvested 48 hours post-transfection by first gently washing them in ice cold phosphate buffered saline (PBS; Sigma-Aldrich), then scraped in ice cold lysis buffer (50 mM Tris pH 7.5, 150 mM NaCl, 10% (v/v) glycerol, 50 mM NaF, 5 mM sodium pyrophosphate, 1% (v/v) Triton X-100, cOmplete protease inhibitor cocktail (Roche)). Lysates were clarified by centrifugation, flash frozen in liquid nitrogen (N_2_) and stored at −80 °C until further analysis.

Overexpressed FLAG-AMPK complexes were immobilised on FLAG-M2 agarose (Sigma-Aldrich A2220) by incubation with lysates for 4 hours at 4 °C. Following gentle centrifugation, the resin was washed twice in high salt purification buffer (50 mM HEPES pH 7.4, 1 M NaCl, 10% (v/v) glycerol, 0.02% (v/v) Tween-20), twice in purification buffer (50 mM HEPES pH 7.4, 150 mM NaCl, 10% (v/v) glycerol, 0.02% (v/v) Tween-20), before being resuspended in FLAG elution buffer (50 mM Tris pH 7.5, 150 mM NaCl, 10% (v/v) glycerol, 0.02% (v/v) Tween-20, 1 mg·mL^−1^ FLAG peptide (Purar Chemical) and 1 mM 1,4-dithiothreitol (DTT)). AMPK was eluted with shaking overnight at 4 °C. Protein concentration was estimated via immunoblotting the FLAG eluate alongside known quantities of bacterially expressed AMPK; the AMPKα antibody was used for all quantification.

For FLAG immunoprecipitation for immunoblot or liquid chromatography mass spectrometry-mass spectrometry (LC-MS/MS) analysis, AMPK was immobilised on FLAG-M2 agarose and washed as above. Following the wash steps, a sample of the resin was taken for immunoblotting and boiled in 3x sample buffer, and the remainder was reserved for LC-MS/MS analysis.

Endogenous ACC was enriched from lysates for immunoblotting using Streptavidin Sepharose High Performance (Cytiva). The resin was incubated with clarified lysates for 2 hours at 4 °C. Following gentle centrifugation, the resin was washed twice in purification buffer before being resuspended and boiled in 3x sample buffer.

### Bacterial protein expression and purification

Recombinant full-length heterotrimeric AMPK _6xHis_α2β2γ1 and _6xHis_α2β2γ3 was expressed in *E. coli* Rosetta 2 (DE3) (Merck Millipore) after double-transformation of pET-Duet-1 (α- and γ-subunits) and pCOLA (β-subunit) plasmids (75). Wild-type and mutant α2-KD (_6xHis-Thrombin_*α*2_6-279_; _6xHis-PreScission_*α*2_6-278_^D139N^; _6xHis-PreScission_*α*2_6-278_^T172D^) was expressed in *E. coli* Rosetta 2 (DE3) after transformation of the pET28a or pET28b plasmid (78). Both heterotrimeric AMPK and kinase domain constructs were purified as described previously (75). Briefly, expression cultures were grown to an optical density (OD_600_) of 0.8 for kinase domain constructs and 3.0 for heterotrimeric AMPK, induced with 500 µM isopropyl-β-D-1-thiogalactopyranoside (IPTG; Gold Biotechnology), followed by overnight incubation at 16 °C with shaking. Cell pellets were resuspended in lysis buffer (50 mM Tris pH 7.6, 500 mM NaCl, 5% (v/v) glycerol, 50 mM imidazole, 2 mM β-mercaptoethanol (BME), 0.01 mM leupeptin, 0.1 mM AEBSF, 0.5 mM benzamidine hydrochloride), lysed using a pre-cooled EmulsiFlex-C5 homogeniser (Avestin) and clarified via centrifugation. Protein was captured on a HisTrap HP 5mL Ni^2+^ column (Cytiva) and eluted with chilled Ni^2+^ column elution buffer (50 mM Tris pH 7.5, 500 mM NaCl, 5% (v/v) glycerol, 400 mM imidazole, 2 mM BME) and separated on a HiLoad 16/600 (Superdex 75 for kinase domain constructs and Superdex 200 for heterotrimeric AMPK) gel filtration column (Cytiva) pre-equilibrated with AMPK size exclusion column buffer (AMPK SEC buffer: 50 mM Tris pH 8.0, 150 mM NaCl, 2 mM Tris(2-carboxyethyl)phosphine (TCEP)). To activate heterotrimeric AMPK, purified protein was incubated with 0.5 mM ATP, 0.5 mM AMP, 2.5 mM MgCl_2_, AMPK SEC buffer and purified CAMKK2 (1:2500 mass ratio of CAMKK2:AMPK; expressed and purified in-house from *E. coli* as previously described (20)) for 1 hour at 22 °C. The phosphorylation reaction was terminated by directly loading onto a HiLoad 16/600 Superdex 200 gel filtration column pre-equilibrated with AMPK SEC buffer. Protein was concentrated to ∼2 mg/mL and ∼10 mg/mL for the kinase domain and heterotrimer complex, respectively, flash frozen in liquid N_2_ and stored at −80 °C.

All recombinant MBP-γ3-NTE and fragments (AVI-tagged and non-AVI-tagged) were expressed in *E. coli* Rosetta 2 (DE3) after transformation of the pMAL-c5X plasmid. Expression cultures were grown at 37 °C in rich broth supplemented with 2 mg·mL^−1^ glucose and 100 µg·mL^−1^ ampicillin to an OD_600_ of 0.6 before induction with 400 µM IPTG and overnight incubation at 16 °C with shaking. To biotinylate the AVI tag, 50 µM biotin was included at the IPTG induction step to enhance *in vivo* biotinylation by endogenous BirA. Cell pellets were resuspended in lysis buffer (50 mM HEPES pH 6.8, 200 mM NaCl, 10 mM BME, 1 mM EDTA, 0.01 mM leupeptin, 0.1 mM AEBSF, 0.5 mM benzamidine hydrochloride), lysed using a pre-cooled EmulsiFlex-C5 homogeniser and clarified via centrifugation. Supernatant was diluted 1:3 with amylose resin buffer (50 mM HEPES pH 6.8, 200 mM NaCl, 10 mM BME, 1 mM EDTA) and passed through amylose resin (NEB) at ∼1 mL·min^−1^. The resin was washed with 10 column volume (CV) of amylose resin buffer and eluted with amylose resin buffer supplemented with 10 mM maltose. Proteinaceous fractions were pooled, concentrated and MBP tag optionally removed by addition of GST-tagged PreScission protease (1:60 w/w, expressed and purified in-house from *E. coli*) incubated overnight at 4 °C. The protein was buffer exchanged into ion exchange buffer A (20 mM Bis-Tris pH 6.0, 20 mM NaCl) using a PD10 desalt column (Cytiva) and loaded onto a Mono Q™ 5/50 GL ion exchange column (Cytiva) pre-equilibrated with ion exchange buffer A. Protein was eluted using a gradient program with ion exchange buffer A and ion exchange buffer B (20 mM Bis-Tris pH 6.0, 1 M NaCl) consisting of 0% B (5 CV), 0-15% B (2 CV), 15% B (10 CV), 15-30% B (40 CV), 30-100% B (10 CV), 100% B (10 CV), 0% B (10 CV). Fractions containing γ3-NTE-AVI or γ3-NTE were pooled and separated on a HiLoad 16/600 (Superdex 75 for MBP-cleaved and Superdex 200 for MBP-tagged) gel filtration column pre-equilibrated with γ3-NTE SEC buffer (20 mM HEPES pH 6.8, 150 mM NaCl, 1 mM maltose, 2 mM TCEP). Fractions containing γ3-NTE-AVI or γ3-NTE were pooled and concentrated to ∼2 mg·mL^−1^, flash frozen in liquid N_2_ and stored at −80 °C. All purified proteins were quality controlled by TOF-MS (Fig. S5A-H), SDS-PAGE Coomassie stain and immunoblotting.

### Radioactive kinase assays

AMPK activity was determined as previously described (70). Briefly, 100 µM SAMS peptide (Purar Chemicals, sequence: HMRSAMSGLHLVKRR), 5 mM MgCl_2_, 200 µM ATP, [γ-^32^P]-ATP (Perkin Elmer), assay buffer (50 mM HEPES pH 7.4, 1 mM DTT and 0.02% (v/v) Tween-20) and purified AMPK were combined and phosphotransferase activity conducted at 30 °C for 10 minutes. Reactions were quenched by spotting 15 µL onto SVI-P cation-exchange paper (79, 80). For determination of the ATP *K*_M_, the concentration of ATP was varied, and all other components were held constant. The papers were repeatedly washed in 1% H_3_PO_4_ (Merck Millipore) and added to vials containing 5 mL Ultima Gold liquid scintillation fluid (Perkin Elmer) and the level of ^32^P-transfer to the SAMS peptide was determined using a Tri-Carb 4810TR liquid scintillation counter (Perkin Elmer). MSG011 was prepared as previously described (36). AMP (A1752) and ATP (A2383) were from Sigma-Aldrich.

### In vitro co-pulldown

Purified _6xHis_α2β2γ1 or _6xHis-PreScission_*α*2_6-278_^T172D^ (or α2-KD(T172D)) was diluted to 10 ng·µL^−1^ in pulldown buffer (50 mM HEPES pH 7.4, 150 mM NaCl, 10% (v/v) glycerol, 0.02% (v/v) Tween-20) and immobilised on Ni^2+^-nitrilotriacetic acid (NTA) Sepharose 6 Fast Flow (Cytiva) by rotating for 2 hours at 4 °C. Following gentle centrifugation, the resin was washed twice in pulldown buffer before being resuspended to a 50% slurry in pulldown buffer. The pulldown was performed in a 100 µL reaction volume containing 25 µL immobilised AMPK (50% slurry in pulldown buffer) and 250 ng γ3_2-168-AVI_ (or γ3-NTE). Other molecules, compounds or peptides were included as indicated, and Ni^2+^-NTA without immobilised _6xHis_α2β2γ1 or _6xHis-PreScission_*α*2_6-278_^T172D^ was included as a negative control. The reactions were incubated at 22 °C with shaking for 1 hour and terminated by gentle centrifugation, removal of the supernatant and washing the resin three times in pulldown buffer. After the final wash the supernatant was removed, 7 µL of Laemmli sample buffer was added and samples boiled and analysed by immunoblotting. The reciprocal pulldown experiment was conducted by performing the method described above, except biotinylated γ3_2-168-AVI_ was immobilised on Streptavidin Sepharose High Performance (Cytiva) and 250 ng of α2-KD(T172D) was added to the assay. SBI-0206965 (SML1540), compound C (P5499), AMP (A1752), ADP (A2754) and ATP (A2383) were from Sigma-Aldrich. Staurosporine (HY-15141) was from MedChemExpress.

### Surface plasmon resonance

SPR single-cycle kinetics were carried out using a Biacore T200 instrument (Cytiva) equipped with an NTA-derivatised carboxymethyldextran sensor chip (NiD200M, XanTec Bioanalytics). All experiments were performed at 25 °C, where both analyte and ligand were diluted in the SPR buffer (50 mM HEPES pH 7.4, 500 mM NaCl, 10% (v/v) glycerol, 50 µM EDTA, 0.05% (v/v) Tween-20). The flow cells were conditioned with 350 mM EDTA (30 µL·min^−1^, 1 min) and activated with 500 µM NiCl_2_ (10 µL·min^−1^, 1 min) before either _6xHis_α2β2γ1 or _6xHis-PreScission_*α*2 ^T172D^ at 20 nM were captured (30 µL·min^−1^, 1 min) at a density of ∼100 RU at the beginning of each cycle. There were two cycles performed for each replicate. A reference cell with neither enhancement nor captured AMPK was used as a blank. To measure the protein-protein interaction, various concentrations (12 nM - 333 nM) of γ3_2-168-AVI_ were injected (30 µL·min^−1^, 1.5 mins). Dissociation was monitored by injection of SPR buffer (30 µL·min^−1^, 5 mins). Sensorgrams were double referenced against the reference cell and buffer reference signal (blank). Affinity (*K*_D_), association rate (*k*_a_), and dissociation rates (*k*_d_) were calculated by global fitting of the data to a 1:1 binding model using Biacore T200 Evaluation Software v2.0.

### Crosslinking

Variants of MBP-tagged AMPK γ3-NTE-AVI (γ3_2-168_, γ3_26-168_, γ3_60-168_, γ3_95-168_, γ3_130-168_) and non-AVI (γ3_1-168_, γ3_2-129_) were bacterially expressed and purified as outlined above in the MBP-tagged format. The MBP tag on the γ3-NTEs was then cleaved by incubating the proteins with PreScission protease (in-house; 1:1000 ratio) overnight. The MBP-cleaved and uncleaved MBP-tagged NTEs were then incubated at 22 °C with either wild-type _6xHis-Thrombin_α2_6-279_ or inactive _6xHis-PreScission_α2_6-278_^D139N^ (or α2-KD(D139N)) (150 r.p.m., 30 min) before 30 mM 4-(4,6-dimethoxy-1,3,5-triazin-2-yl)-4-methylmorpholinium chloride (DMTMM, Sigma-Aldrich 74104-1G-F) or 50 mM HEPES pH 7.5 (negative control) was spiked in for 30 min at 22 °C. The reactions were quenched with the addition of sample buffer then separated by SDS-PAGE, followed by immunoblot.

The MBP-cleaved γ3_2-168_ and _6xHis-Thrombin_α2_6-279_ were also crosslinked with sulfosuccinimidyl-diazirine (sulfo-SDA, Thermo Scientific Pierce 26173) using a protein-to-crosslinker ratio of 1:0.5 (w/w). All materials were dissolved in crosslinking buffer (50 mM HEPES pH 7.5, 20 mM NaCl, 10% (v/v) glycerol). The reactions were incubated for 60 min at 22 °C in the dark. The diazirine group was then photoactivated by ultraviolet light radiation using a UVP CL-1000L UV cross-linker. Samples were prepared as thin films on Eppendorf tube lids, placed on ice 5 cm from the ultraviolet light, and irradiated for 1 minute at 365 nm before quenching the reaction with 1 M Tris pH 7.5 at a 1:15 ratio. Post-irradiation, the crosslinked samples were separated via SDS-PAGE.

### HDX-MS

HDX labelling of protein was performed at 20 °C for periods of 0, 6, 60, 600 and 6000 seconds using a PAL Dual Head HDX Automation manager (Trajan/LEAP) controlled by the ChronosHDX software. 3 µL of 15 µM _6xHis-Thrombin_α2_6-279_ or γ3_2-168-AVI_ or a 1:1 mixture of both proteins (7.5 µM of each protein incubated for 30 min at 22 °C) was transferred to 55 µL of non-deuterated (5 mM potassium phosphate buffer pH 7.0 in H_2_O) or deuterated (5 mM potassium phosphate buffer pD 7.0 in D_2_O) buffer and incubated for the respective time. Labelling was quenched by adding 50 µL of the deuterated protein to 50 µL of quench buffer (50 mM potassium phosphate buffer pH 2.3) at 1 °C. For online pepsin digestion, 80 µL of the quenched sample was passed over an immobilised 2.1 × 30 mm Enzymate BEH pepsin column (Waters) equilibrated in 0.1% formic acid in H_2_O at 100 µL/min. To further reduce peptide carryover, n-Octyl-β-d-glucopyranoside at 1% w/w was added the pepsin column wash solution (1.5 M guanidine hydrochloride, 4% acetonitrile, 0.8% formic acid). Proteolyzed peptides were captured and desalted by a C18 trap column (VanGuard BEH; 1.7 μm; 2.1 × 5 mm; Waters) and eluted with acetonitrile and 0.1% formic acid gradient (5% to 40% in 8 min, 40% to 95% in 0.5 min, 95% 1.5 min) at a flow rate of 80 μL/min and separated on an ACQUITY UPLC BEH C18 analytical column (1.7 μm, 1 × 50 mm, Waters) delivered by ACQUITY UPLC I-Class Binary Solvent Manager (Waters).

For mass spectrometry, an ion mobility equipped SYNAPT G2-Si mass spectrometer (Waters) was used. Instrument settings were: 3.0 KV capillary and 40 V sampling cone with source and desolvation temperature of 100 °C and 40 °C respectively. The desolvation and cone gas flow was at 800 L/hr and 100 L/hr, respectively. High energy ramp trap collision energy was from 20 V to 40 V. All mass spectra were acquired using a 0.4 sec scan time with continuous lock mass (Leu-Enk, 556.2771 m/z) for mass accuracy correction. Data were acquired in HDMS^E^ (ion mobility) mode and peptides from non-deuterated samples were identified using Protein Lynx Global Server (PLGS) v3.0 (Waters). To ensure high peptide selection stringency, we applied additional filter constraints of 0.3 fragments per residue, minimum intensity of 5000, maximum [M+H]^+^ error of 5 ppm, retention time RSD of 10% and file threshold of 4 out of 6 HDMS^E^ files. The deuterium uptake values were calculated for each peptide using DynamX 3.0 (Waters). The average back-exchange in our system (∼30%) was measured using phosphorylase B, but we did not adjust for back exchange as our analysis compares relative deuteration between protein states, and therefore all results are reported as relative deuterium exchange levels expressed in mass units (Da). Deuterium exchange experiments were performed in triplicate for each of the timepoints. Peptides with a statistically significance difference in HDX were determined using the Deuteros 2.0 software with a hybrid Woods differential significance test with a 99% confidence interval.

### Structure prediction of γ3-AMPK with AlphaFold Server

Using AlphaFold3 server, the γ3-NTE interaction with α2-KD were modelled from their protein sequences individually in isolation and as part of AMPK heterotrimer complex. For modelling accuracy, the AMPK sequences were adapted to integrate our current experimental findings from complementary biophysical techniques including HDX-MS, SPR, chemical crosslinking and co-pulldowns (Table S1). The prediction confidence level was determined from its integrated AlphaFold confidence metrics which are predicted local distance different test (pLDDT; 0 to 100, with 100 indicates the highest confidence) shown as colour outputs in the image structure and predicted template modelling (pTM; 0 to 1, with 1 indicates the overall predicted structure being the most similar to the experimental published structure). Images of the predicted structures with the highest confidence level were viewed and created on PyMOL.

### LC-MS analysis on peptides

FLAG immobilised AMPK for peptide analysis was prepared as previously described (46). Briefly, protein was precipitated on the FLAG resin by addition of 100% methanol and incubated on ice for 30 mins. The sample was centrifuged, supernatant discarded, and resin dried under N_2_ gas. The dried resin was resuspended with 50 mM Tris pH 7.5 and 200 ng of Trypsin (Promega) was added, the samples were digested overnight shaking at 37 °C. All digests were taken for LC-MS/MS analysis.

All peptide analysis was carried out on a TripleTOF 5600 mass spectrometer (Sciex) operated with the turbo V DuoSpray ion source linked to an Ultimate 3000 RSLCnano system loading pump (Dionex) and Ultimate 3000 RS autosampler (Dionex). The LC-MS was operated using the Analyst TF v1.7.1 software (Sciex). Source and collision gas was provided by a Genius NM3G nitrogen gas generator (PEAK Scientific). Peptides were resolved on a Waters Acquity BEH peptide C18 column (100 mm x 2.1 mm, 1.7 µm, 130 Å) using an LC solvent system comprised of H_2_O with 0.1% formic acid for channel A, and 90% acetonitrile/10% H_2_O with 0.1% formic acid for channel B. The mass spectrometer was set to time of flight-mass spectrometry (TOF-MS) acquisition mode and operated in positive ion mode with a mass range of 200-3000 m/z and accumulation time of 0.5 seconds. TOF-MS/MS data was visualized using PeakView v2.2 software (Sciex) and peptide peak areas quantified using Skyline v21.1 (MacCross Lab Software). Phosphorylation stoichiometries were calculated from phosphorylated and dephosphorylated peptide peak areas as previously described (46).

### Statistical analysis

All statistical analyses were performed using Prism v10.6.1(GraphPad Software). Results from replicate experiments (*n*) are expressed as means −/+ standard error (s.e.m.) unless stated otherwise.

## Data availability

All data are contained within the manuscript and available upon request. AlphaFold3 structure prediction data have been deposited in figshare (81).

## Supporting information

This article contains supporting information.

## Acknowledgements

S.G., J.W.S., B.E.K. and J.S.O. were supported by National Health and Medical Research Council (NHMRC) project grants (GNT1145836, GNT1138102 and GNT2029423). K.S. was supported from the Novo Nordisk Foundation (NNF23SA0084103). C.R.H. was supported by an NHRMC early career research fellowship (GNT2034044). C.G.L. was supported by the Jack Brockhoff Foundation (no. JBF-4206), EH Flack fellowship, and NHMRC early career research fellowship (GNT1143080). This project was supported by St Vincent’s Institute of Medical Research (Australia) and in part by the Victorian Government’s Operational Infrastructure Support Program. We would like to thank Yi Sing Gee for the kind donation of the compound MSG011. We acknowledge the use of the Melbourne Mass Spectrometry and Proteomics Facility at the Bio21 Molecular Science and Biotechnology Institute.

## Author contributions

A.J.O., M.N.H.K., D.Y., N.X.Y.L., A.H., D.A.O., A.C.P.G, M.C., G.X.Y.Z, B.R.T., L.D.,

C.S.A. and C.R.H. provided reagents and performed the experiments. W.J.S., J.W.S., K.S., M.W.P., B.E.K. and S.G. provided conceptual input. C.G.L. and J.S.O. conceived the study. A.J.O., M.N.H.K., J.W.S., J.S.O. and C.G.L. wrote the initial draft and all authors contributed to the final manuscript.

## Supplementary Figures

**Figure S1.**
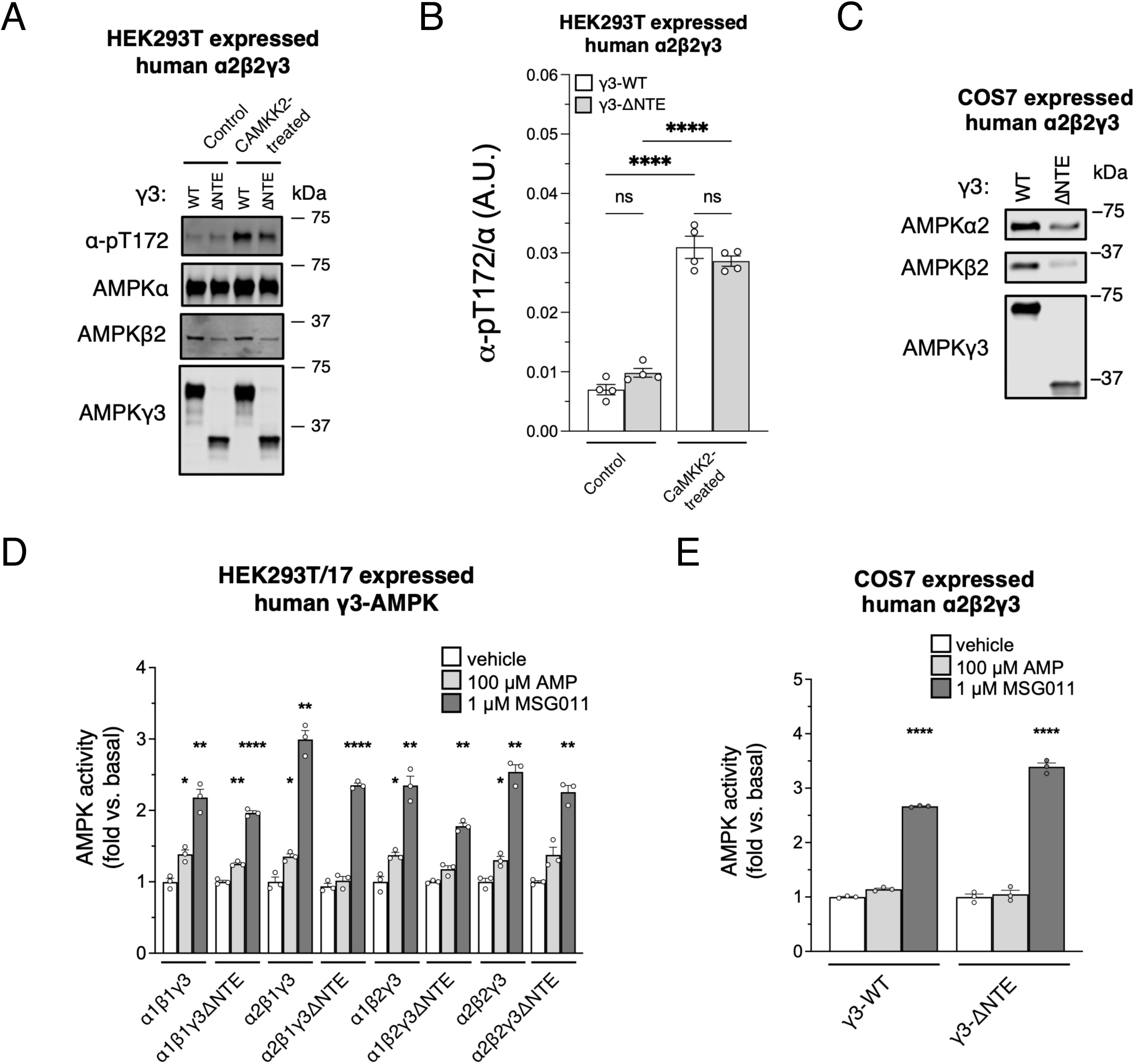
Validation of AMPK complexes overexpressed in cultured mammalian cells. Related to Figure 1. (A) Immunoblot images of α-pT172 and AMPK subunits from α2β2- complexes containing γ3-WT or γ3-ΔNTE, expressed and FLAG-purified (with or without CAMKK2-treatment) from HEK293T cells. (B) Densitometry analysis of (A) comparing basal α-pT172 levels between γ3-complexes. Data are shown as mean α-pT172 level ± s.e.m.; *n* = 4. Statistical analyses were performed by one-way ANOVA with Dunnett’s multiple-comparisons test vs. γ3-WT, \*\*\*\**p* < 0.0001, or CAMKK2-untreated, *ns* > 0.05. Representative immunoblot images of 4 independent experiments are shown. (C) Immunoblot images of AMPK α2β2-complexes containing γ3-WT or γ3-ΔNTE, FLAG-purified from COS7 cells. (D) *In-vitro* AMP- or MSG011-stimulated activities of γ3-WT and γ3-ΔNTE AMPK complexes from HEK293T/17 cells, expressed as fold-change relative to vehicle treated. Data are shown as mean fold-change in AMPK activity ± s.e.m.; *n* = 3. Statistical analyses were performed by one-way ANOVA with Dunnett’s multiple-comparisons test vs. vehicle treated. \**p* < 0.05, \*\**p* < 0.01, \*\*\*\**p* < 0.0001. (E) AMP- or MSG011-stimulated activities of AMPK complexes from (C), expressed as fold-change relative to vehicle treated. Data are shown as mean fold-change in AMPK activity ± s.e.m.; *n* = 3. Statistical analyses were performed by one-way ANOVA with Dunnett’s multiple-comparisons test vs. vehicle treated. \*\*\*\**p* < 0.0001.

**Figure S2.**
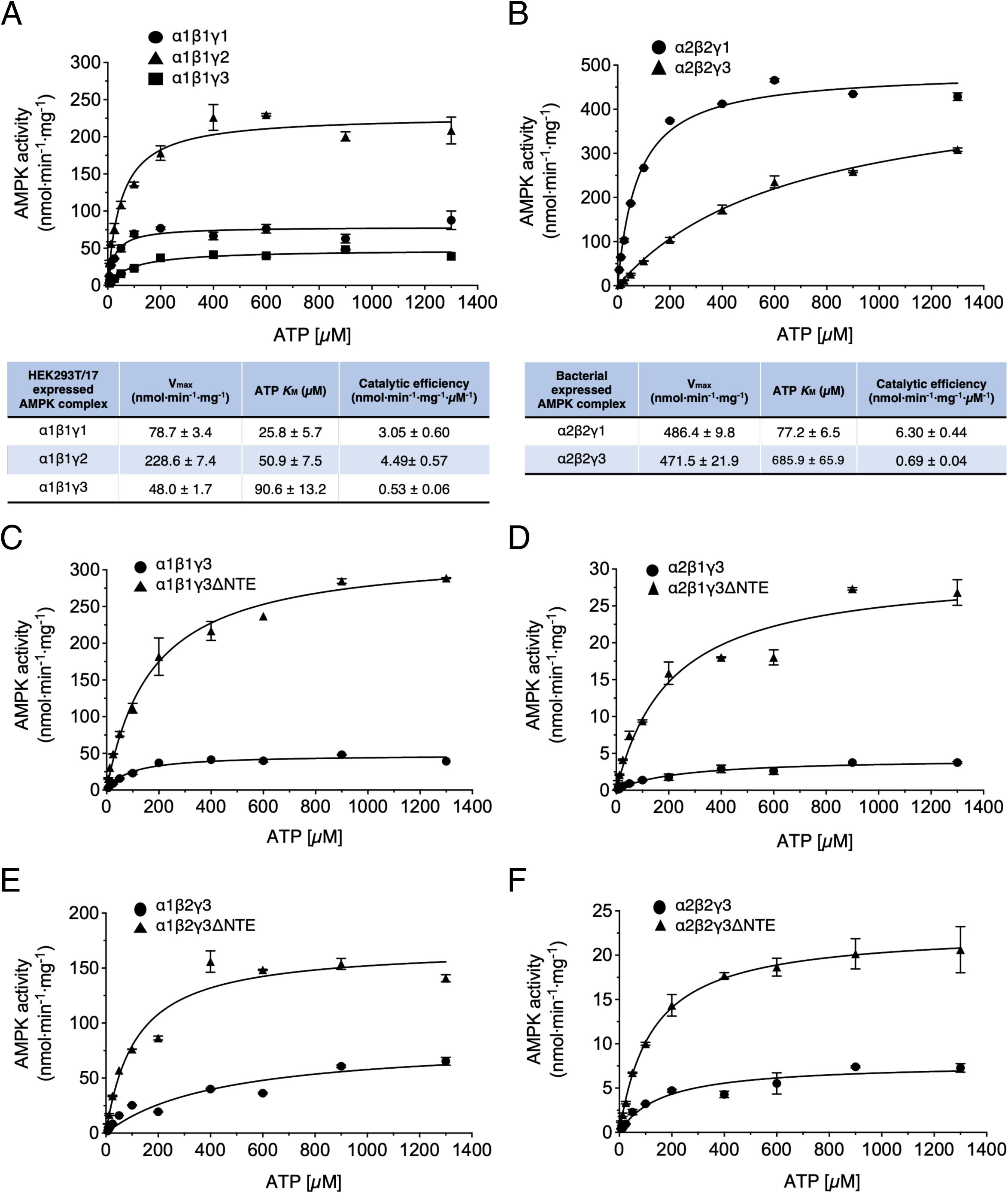
Kinetics of AMPK activity with γ3-WT and γ3-ΔNTE. Related to Table 1. ATP dose-response curves for (A) AMPK α1β1γ1, α1β1γ2 and α1β1γ3, FLAG-purified from HEK293T/17 cells, and (B) AMPK α2β2γ1 and α2β2γ3 purified from *E. coli*. Kinetic parameters are detailed in the associated table. These values were derived from Michaelis-Menten non-linear fit regression with each parameter presented as value ± s.e.m. (C-F) ATP dose-response curves for AMPK containing γ3-WT or γ3-ΔNTE, FLAG-purified from HEK293T/17 cells, providing kinetics data presented in Table 1. All data are presented as mean AMPK activity ± s.e.m., *n* = 2.

**Figure S3.**
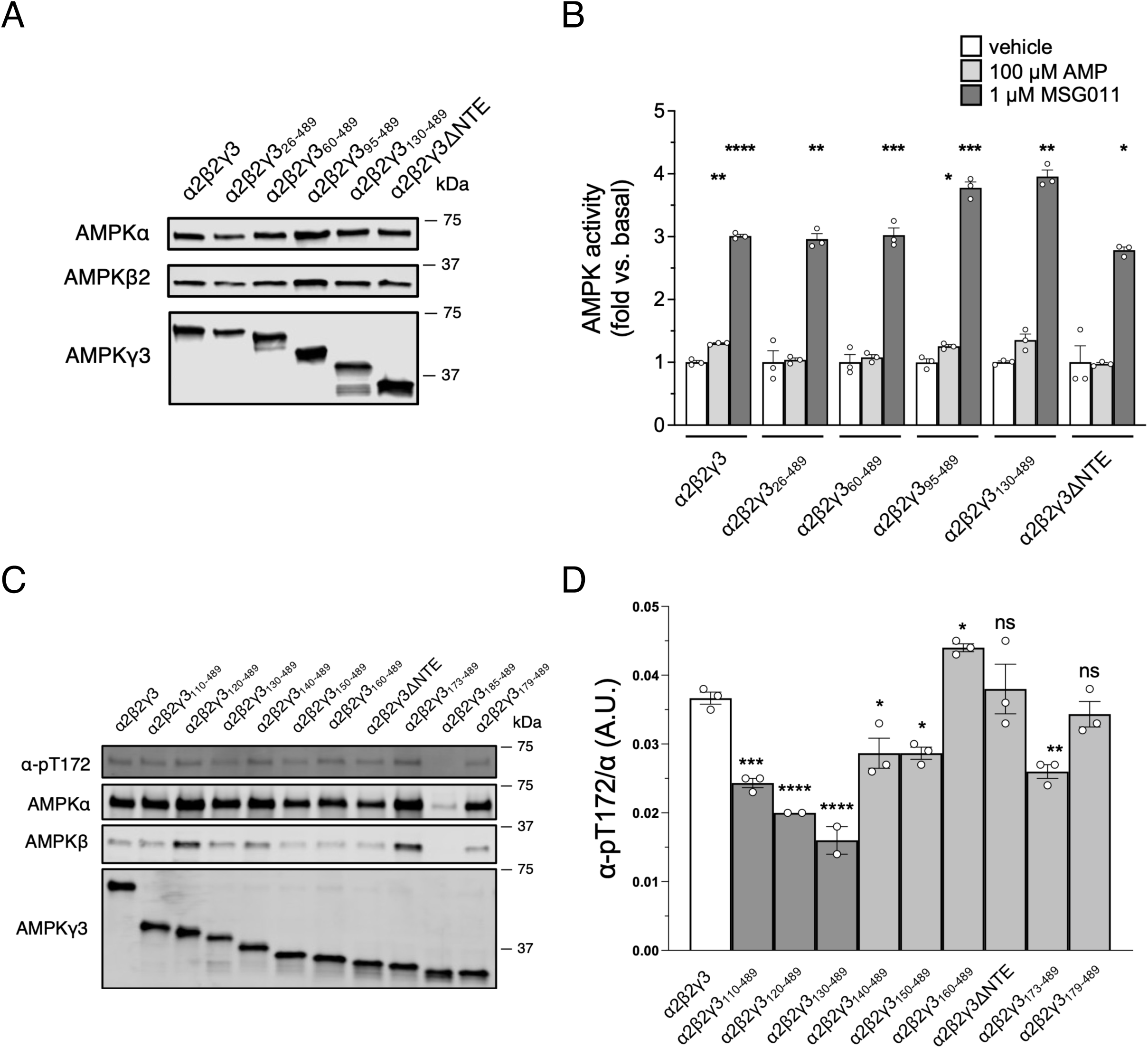
Expression and activities of AMPK α2β2-complexes containing various γ3- NTE truncations. Related to Figure 1. (A) Immunoblot images of AMPK α2β2-complexes containing γ3-WT or sequential γ3 truncations, expressed and FLAG-purified from HEK293T/17 cells. (B) *In-vitro* AMP- or MSG011-stimulated activities of AMPK complexes from (A), expressed as fold-change relative to vehicle treated. Data are shown as mean fold-change in AMPK activity ± s.e.m.; *n* = 3. Statistical analyses were performed by one-way ANOVA with Dunnett’s multiple-comparisons test vs. vehicle treated. \**p* < 0.05, \*\**p* < 0.01, \*\*\**p* < 0.001, \*\*\*\**p* < 0.0001 vs. vehicle treated. (C) Immunoblot images of α-pT172 and AMPK subunits from α2β2- complexes containing γ3-WT or finer sequential γ3 truncations, expressed and FLAG-purified from HEK293T cells. (D) Densitometry analysis of (C) comparing basal α-pT172 between the complexes. Data are shown as mean α-pT172/α (A.U.) ± s.e.m.; *n* = 2-3. Statistical analyses were performed by by one-way ANOVA with Dunnett’s multiple-comparisons test vs. γ3-WT. *ns* > 0.05, \**p* < 0.05, \*\**p* < 0.01, \*\*\**p* < 0.001, \*\*\*\**p* < 0.0001. Representative immunoblot images of 3 independent experiments are shown.

**Figure S4.**
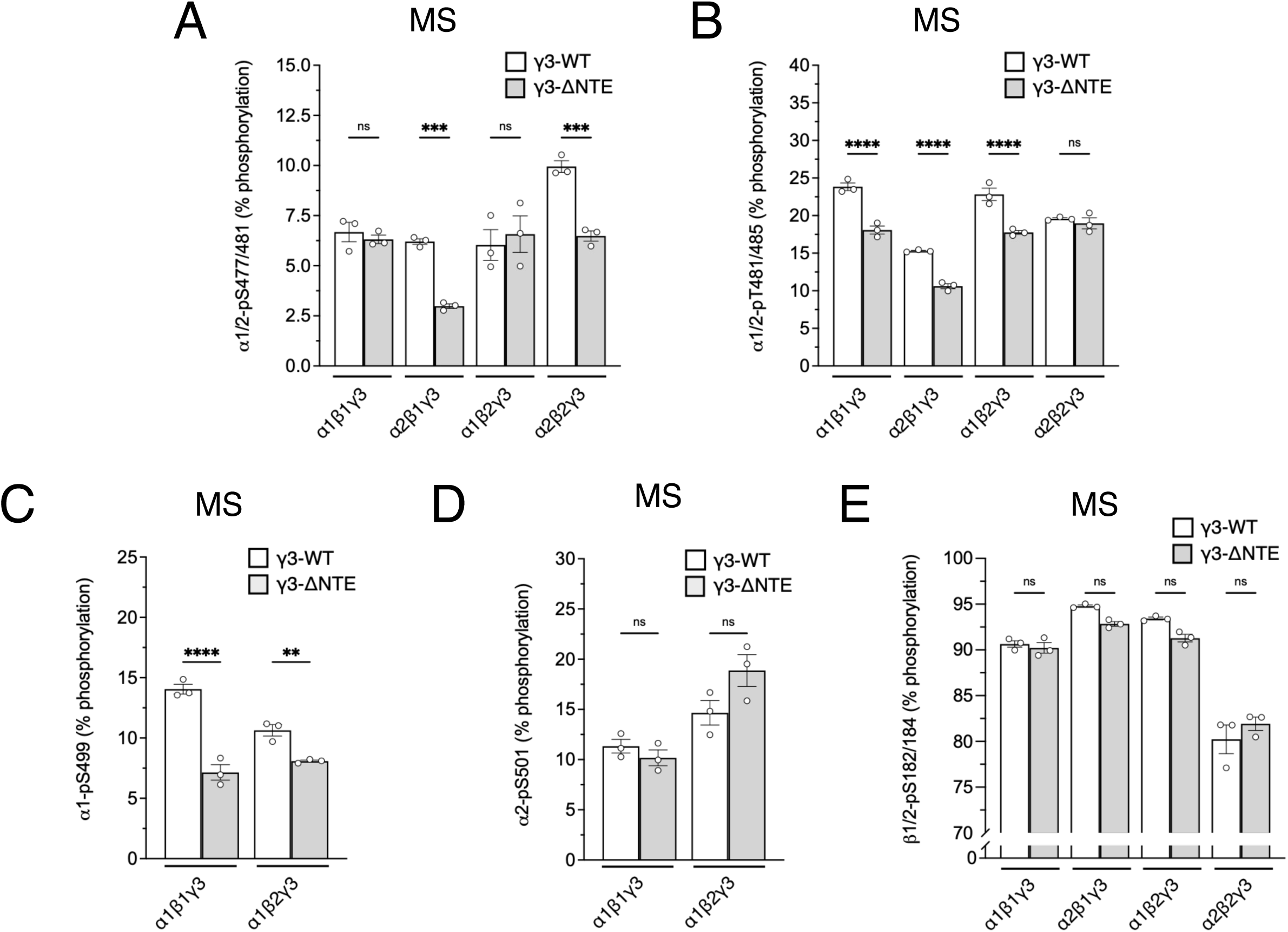
Phosphorylation site profiling of AMPK complexes with γ3-WT or γ3-ΔNTE. Related to Figure 2. (A-E) Area under the curve stoichiometry analysis of LC-MS data for tryptic-digested from the FLAG-purified AMPK for multiple phosphorylation sites (A) α1/2- pS477/481, (B) α1/2-pT481/485, (C) α1-pS499, (D) α2-p S501, and (E) β1/2-pS182/184. Data are adjusted for the flyability ratios as previously mentioned (46), and presented as mean % phosphorylation ± s.e.m.; *n* = 3. Statistical analyses were performed by Welch’s *t*-test vs. respective γ3-WT complexes. *ns*, *p* > 0.05, \*\**p* < 0.01, \*\*\**p* < 0.001, ****p < 0.0001.

**Figure S5.**
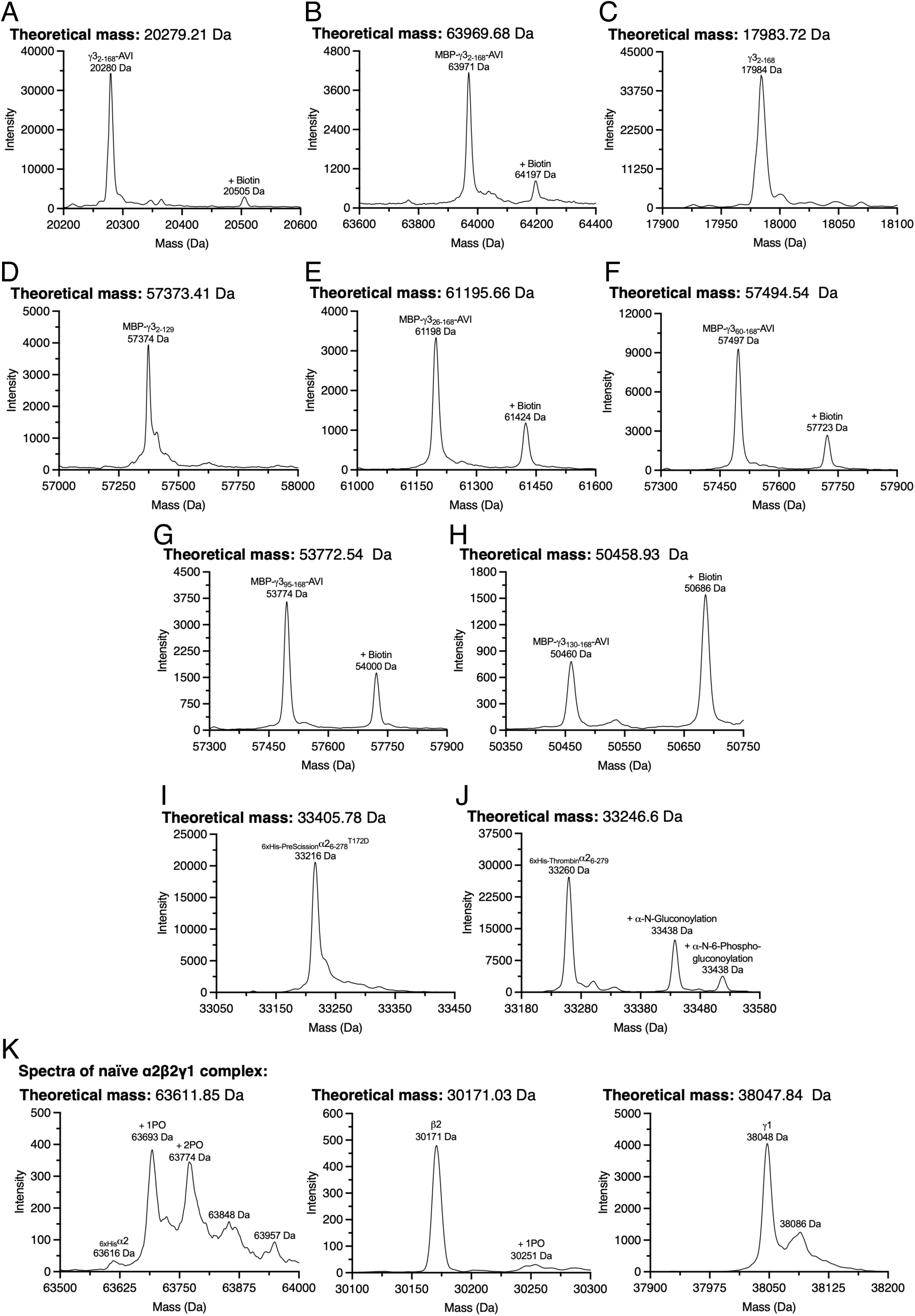
Reconstructed intact mass spectra of bacterially expressed protein. Reconstructed, intact protein mass spectra confirming the molecular weight and their respective post-translational modification status of purified: (A) γ3_2-168-AVI_; (B) MBP-γ3_2-168- AVI_; (C) γ3_2-168_; (D) MBP-γ3_2-129_; (E) MBP-γ3_26-168-AVI_; (F) MBP-γ3_60-168-AVI_; (G) MBP-γ3_95-168-AVI_; (H) MBP-γ3_130-168-AVI_; (I) _6xHis-PreScission_α2_6-278_^T172D^; (J) _6xHis-Thrombin_α2_6-279_; (K) naïve_6xHis_α2β2γ1.

**Figure S6.**
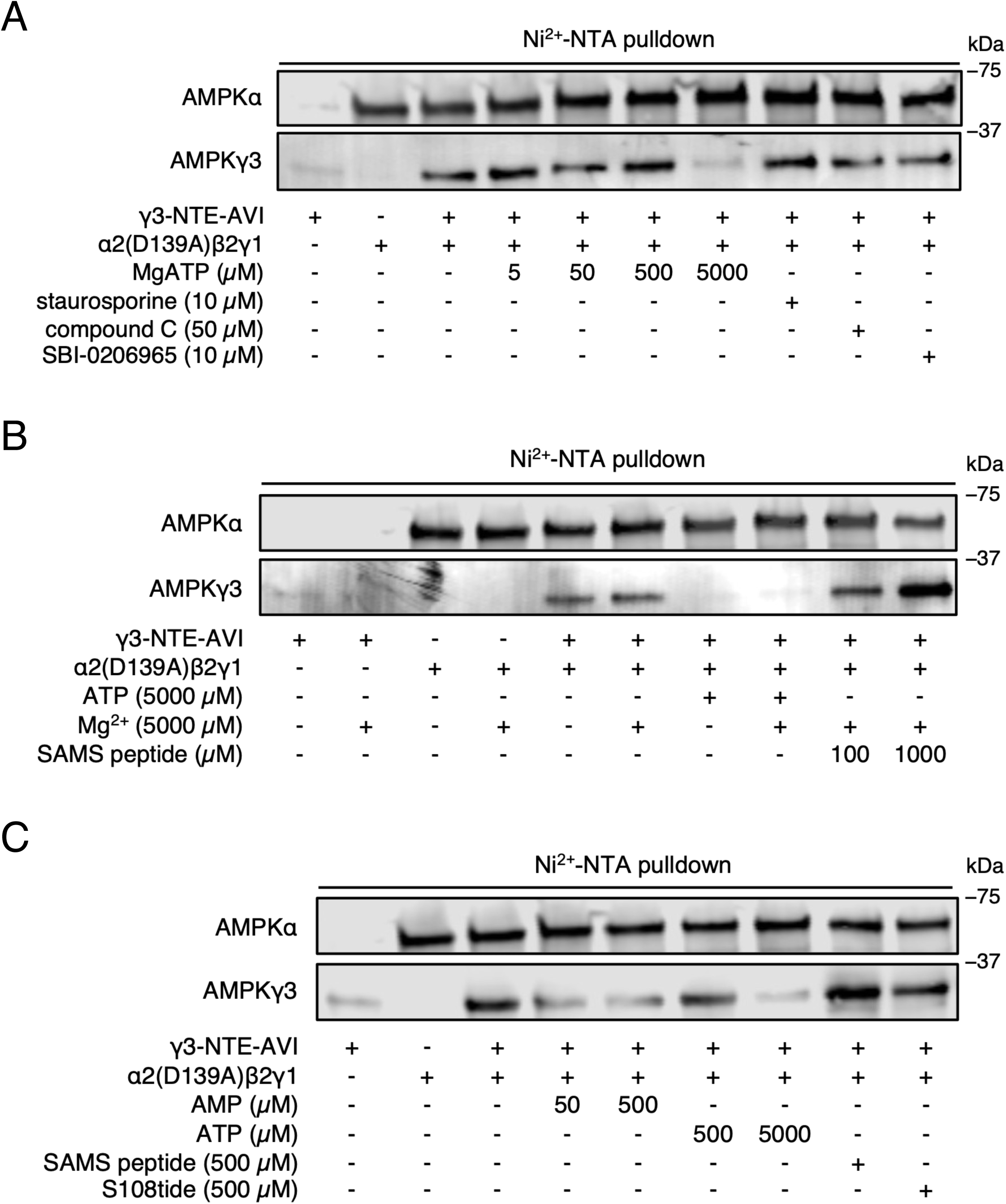
AMPK γ3-NTE directly interacts with the AMPK heterotrimer. Related to Figure 2. Immunoblot images of co-pulldown experiments using Ni^2+^-NTA- immobilised α2(D139A)β2γ1 heterotrimer (kinase dead) to capture soluble γ3-NTE, individually pre-incubated with (A) Mg^2+^, ATP or inhibitors staurosporine, compound C or SBI-0206965, (B) ATP, Mg^2+^ or SAMS peptide, or (C) AMP, ATP, SAMS peptide or S108tide. Bound material was immunoblotted for AMPKα and γ3. Representative immunoblots from 3 independent experiments are shown.

**Figure S7.**
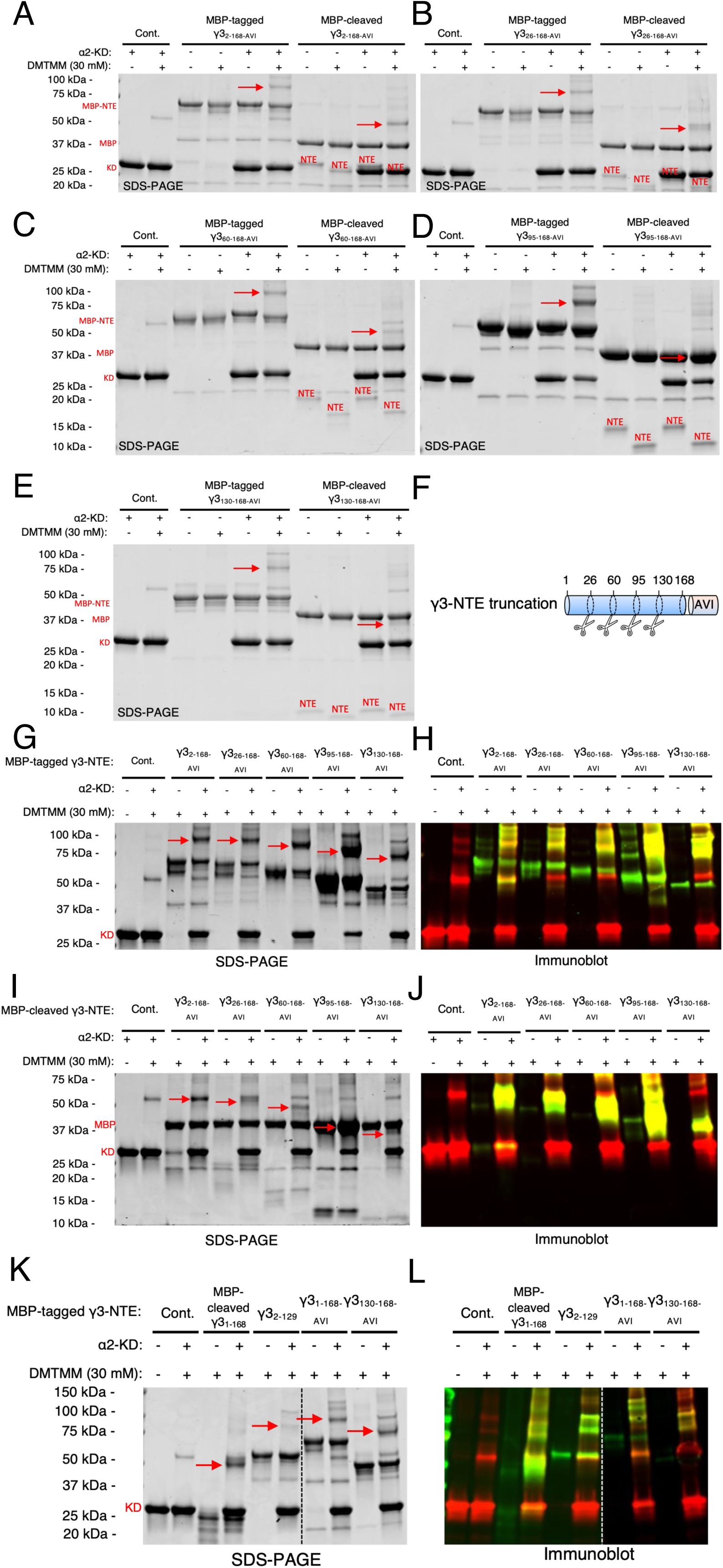
DMTMM-mediated crosslinking of AMPK α2-KD with MBP-tagged/cleaved γ3-NTE truncations. Related to Figure 4. SDS-PAGE images of DMTMM crosslinking reactions using α2-KD and MBP-tagged/cleaved (A) γ3_2-168_, (B) γ3_26-168_, (C) γ3_60-168_, (D) γ3_95-168_, and (E) γ3_130-168_. MBP- γ3-NTE, γ3-NTE, cleaved MBP and α2-KD bands are also indicated. (F) Visualization of γ3- NTE truncations used in this figure. (G) SDS-PAGE and (H) corresponding immunoblot images of DMTMM crosslinking reactions using α2-KD and MBP-tagged, sequential γ3-NTE truncations. (I) SDS-PAGE and (J) corresponding immunoblot images of DMTMM crosslinking reactions using α2-KD and MBP-cleaved, sequential γ3-NTE truncations. (K) SDS-PAGE and (L) corresponding immunoblot images of DMTMM crosslinking reactions using α2-KD and various γ3-NTE fragments. For SDS-PAGE images, red arrows indicate α2- KD/γ3-NTE crosslinked species. In some cases, α2-KD/γ3-NTE crosslinked species co- migrated with α2-KD crosslinked pairs or with cleaved MBP. For immunoblot images, red bands = α2-KD (AMPKα antibody), green bands = γ3-NTE (AVI-tag or AMPKγ3 antibody); yellow (merged) bands = α2-KD/γ3-NTE crosslinked species.

**Figure S8.**
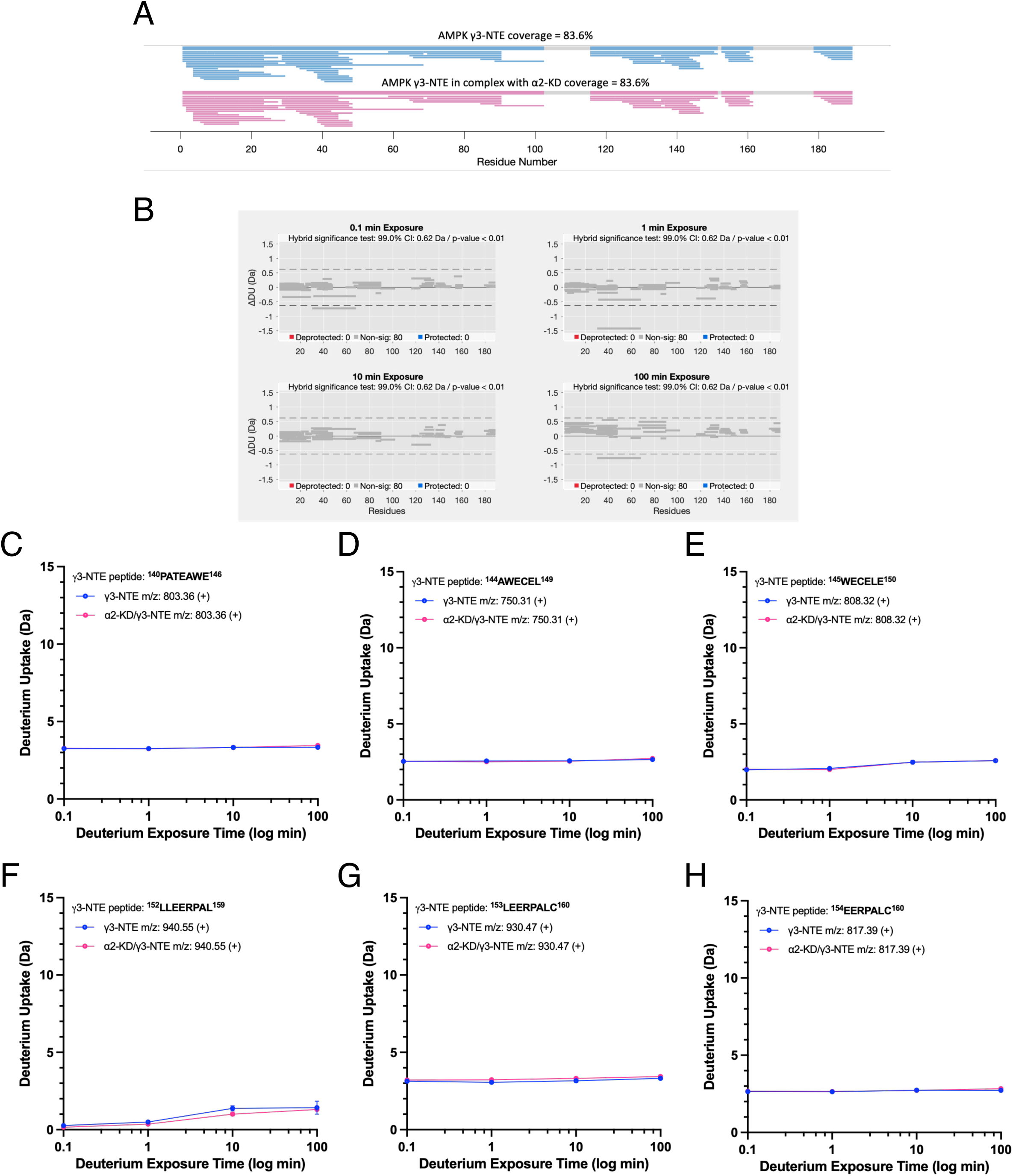
HDX-MS data for AMPK γ3-NTE in the presence of α2-KD. Related to Figure 5. (A) Visualisation of the HDX-MS protein sequence coverage for γ3-NTE. (B) Woods plots for differences in deuterium uptake (ΔDU) by γ3-NTE ± α2-KD across the three time points of deuterium exposure; 0.1 min, 1 min, 10 min and 100 min. A hybrid significance test consisting of a two-prong statistical test, implemented in Deuteros 2.0 was used to identify peptides that show a significant difference. Dashed horizontal lines indicate 99% confidence limit applied to the dataset to identify peptides with significant deuteration differences. Grey = unchanged peptides. No significantly protected or deprotected peptides were detected. (C-H) Deuterium uptake plots over deuterium exposure time for γ3- NTE peptides detected in HDX-MS experiments covering residues within γ3_140-160_.

**Figure S9.**
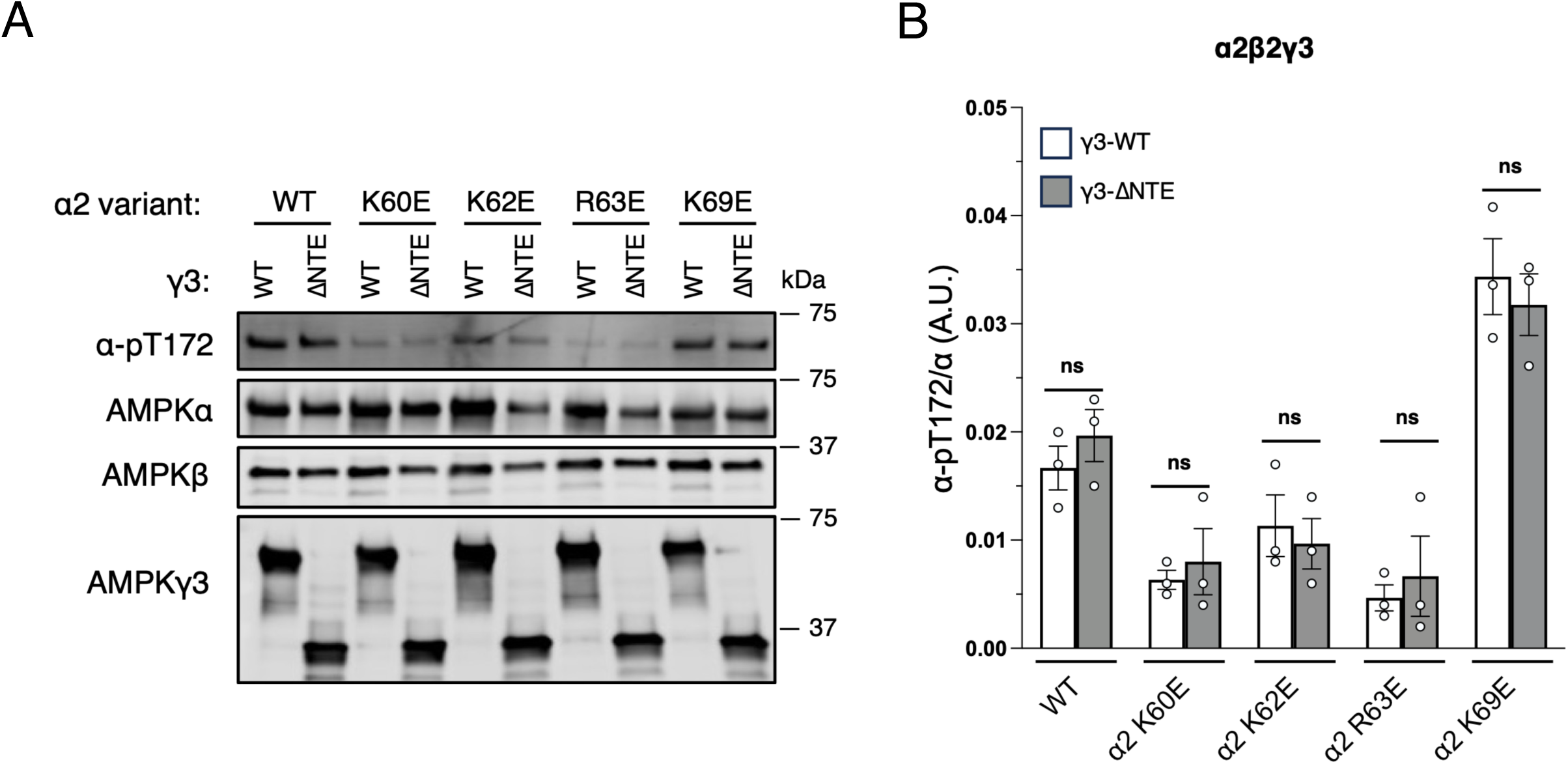
Expression, pT172 levels and basal activities of AMPK α2β2-complexes containing α2 mutations. Related to Figure 5. (A) Immunoblot images of α-pT172 and AMPK subunits from complexes containing α2-WT or mutations in combination with β2 and γ3-WT or γ3-ΔNTE, expressed and FLAG-purified from HEK293T cells. (B) Densitometry analysis of (A) comparing basal α-pT172 levels of complexes containing γ3-ΔNTE to their respective γ3-WT. Data are shown as mean α-pT172 /α (A.U.) ± s.e.m.; *n* = 3. Statistical analyses were performed by Welch’s *t*- test vs. respective γ3-WT complexes. *ns*, *p* > 0.05.

**Table S1.**
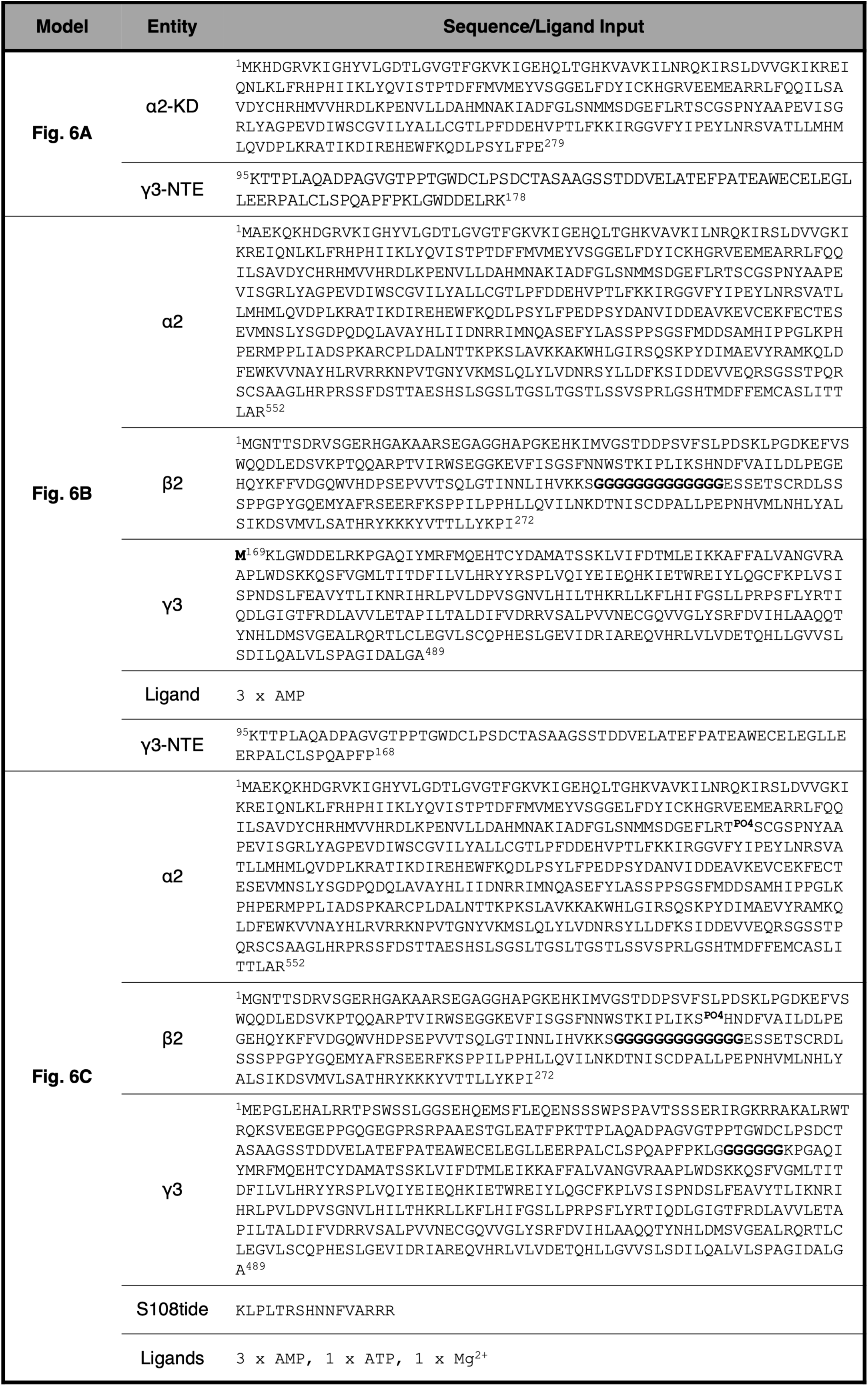
Related to Figure 6. AlphaFold input for γ3-AMPK structural prediction.

**Figure S10.**
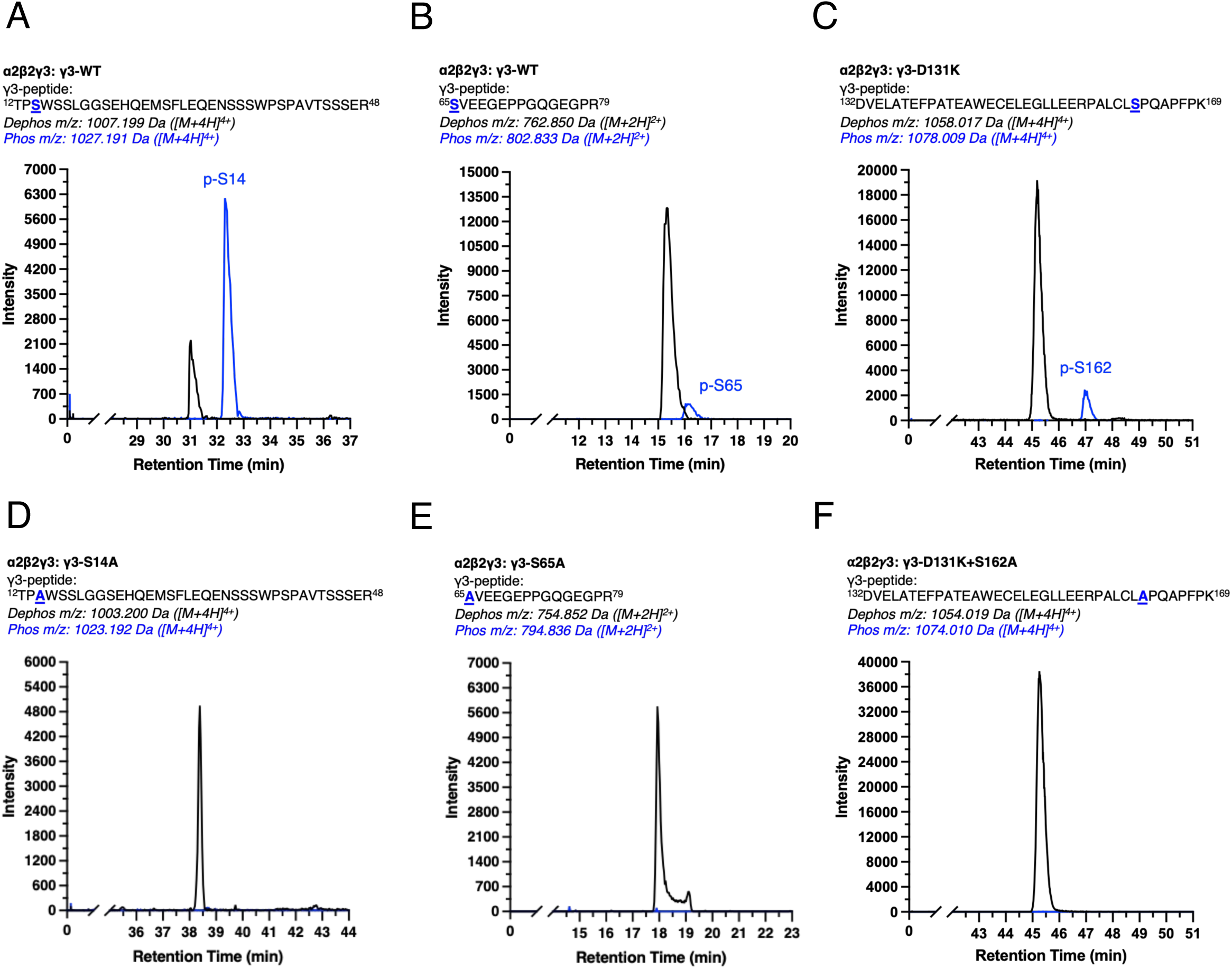
Representative MS-extracted ion counts of phosphorylated residues on the γ3-NTE. AMPK α2β2γ3 (γ3 WT or mutants as indicated) was FLAG-purified from HEK293T cells, digested with trypsin and analysed by LC-MS/MS. Example extracted ion counts (EIC) for cognate phosphorylated and dephosphorylated peptides for (A) γ3_12-48_, (B) γ3_65-79_, (C) γ3_132-169_ with a γ3-D131K mutation, (D) γ3_12-48_ with a γ3-S14A mutation, (E) γ3_65-79_ with a γ3-S65A mutation, and (F) γ3_132-169_ with a γ3-D131K+S162A double mutation.

